# Structure and mechanism of human sphingosine-1-phosphate transporter MFSD2B

**DOI:** 10.64898/2026.01.16.700024

**Authors:** Shahbaz Ahmed, Min Huang, Yaxin Dai, Xuebo Yang, Chia-Hsueh Lee, Long N. Nguyen

**Affiliations:** Department of Structural Biology, St. Jude Children’s Research Hospital, Memphis, TN, USA; Department of Biochemistry, Yong Loo Lin School of Medicine, National University of Singapore, Singapore; Key Laboratory of Structure-Based Drugs Design & Discovery of Ministry of Education, Shenyang Pharmaceutical University, Shenyang 110016, China; Immunology Program, Life Sciences Institute, National University of Singapore, Singapore 117456; Singapore Lipidomics Incubator (SLING), Life Sciences Institute, National University of Singapore, Singapore 117456; Cardiovascular Disease Research (CVD) Program, Yong Loo Lin School of Medicine, National University of Singapore, Singapore 117545; Immunology Translational Research Program, Yong Loo Lin School of Medicine, National University of Singapore, Singapore 117456

## Abstract

Sphingosine-1-phosphate (S1P) is an essential signaling lipid that maintains vascular integrity and regulates immune cell trafficking. The major facilitator superfamily domain–containing protein 2B (MFSD2B) serves as the main S1P exporter in red blood cells and platelets; however, its structure and transport mechanism are unclear. Here, we report the 3.0 Å cryo-EM structure of human MFSD2B bound to S1P. S1P is captured in a distinctive binding state, deeply buried within the C-domain, with its sphingoid tail accommodated by a hydrophobic pocket and its phosphate group coordinated by a cluster of polar residues within the transporter’s cavity. Mutagenesis and molecular dynamics simulations identify the TM2/TM11 lateral opening as the primary pathway for S1P translocation, with key charged residues acting as sequential anchors during transport. Furthermore, we demonstrate that MFSD2B functions as a uniporter, and that subtle rewiring of local charge networks can alter its coupling mechanism. Our work provides a molecular framework for understanding S1P transport mediated by MFSD2B in hematopoietic cells.

## Introduction

Sphingosine-1-phosphate (S1P) is a potent bioactive sphingolipid that orchestrates a wide spectrum of physiological processes, including vascular integrity, cell survival, and lymphocyte trafficking^1–5^. In most cells, S1P is synthesized from sphingosine by sphingosine kinases 1 and 2 (SphK1/2)^6–8^. A specialized group of cell types, most notably vascular endothelial (both blood and lymphatic endothelia) and hematopoietic cells, can efficiently export S1P to the extracellular milieu, where it binds to albumin and high-density lipoproteins (HDL) for autocrine or paracrine signaling functions. However, due to its amphipathic nature, S1P cannot freely diffuse across lipid bilayers and requires dedicated transporters to mediate its translocation.

While several ATP-binding cassette transporters, including ABCA1, ABCC1, and ABCG2, have been implicated in S1P efflux^9,10^, only two transporters have been identified as bona fide S1P transporters: protein spinster homolog 2 (SPNS2) and major facilitator superfamily domain–containing 2B (MFSD2B)^11–14^. SPNS2, expressed predominantly in endothelial cells, mediates S1P secretion into lymph and is thereby critical for regulation of lymphocyte egress from lymph nodes^15–17^. In contrast, MFSD2B is highly expressed in hematopoietic cells, including red blood cells (RBCs) and platelets, where it drives the release of S1P into the circulation^14^. Mouse studies have revealed that SPNS2 and MFSD2B together account for more than 80% of plasma S1P^12,18–20^. Ablation of these sources of S1P caused hemorrhagic and lethality in mouse embryos, underscoring their central roles in maintaining systemic S1P homeostasis^18^.

We previously identified MFSD2B as the principal S1P exporter in RBCs and platelets using genetic knockout models. Lipidomic profiling demonstrated that deletion of MFSD2B causes S1P accumulation in these cells and reduces circulating S1P levels by ∼54% relative to wild-type mice^14^. Among the known cell types where MFSD2B is expressed, RBCs have been demonstrated as the major source for circulating S1P, while platelets release a small amount of S1P upon activation^21^. Loss of MFSD2B disrupts RBC and platelet morphology and function, highlighting its essential role in blood cells’ physiology^18,21^. MFSD2B has also been implicated in cardiovascular function, although the mechanism is unclear^22^.

MFSD2B belongs to the major facilitator superfamily (MFS). Among MFS transporters, four members, MFSD2A, MFSD2B, SPNS1, and SPNS2, have been characterized as lysolipid transporters. MFSD2B and SPNS2 mediate S1P transport across the plasma membrane, whereas MFSD2A and SPNS1 translocate lysophosphatidylcholine (LPC) and lysophosphatidylethanolamine (LPE) via the plasma membrane and lysosomes, respectively^23–25^. Recent structural studies of MFSD2A, SPNS1, and SPNS2 have revealed how these transporters move amphiphilic substrates across cell membranes^26–35^: MFSD2A imports its substrates in a Na⁺-dependent manner, while SPNS1 exports lysosomal lysolipids utilizing a gradient of protons. In contrast, SPNS2 functions as a uniporter that drives S1P efflux down its concentration gradient. Despite these advances, the molecular mechanism by which MFSD2B recognizes and translocates S1P across the membrane remains unknown.

Here, we present the cryo-EM structure of human MFSD2B bound to S1P at 3.0 Å resolution. The structure captures a distinctive outward-facing conformation in which S1P binds deep within the transporter, coordinated by a hydrophobic pocket that accommodates its sphingoid tail and a hydrophilic cavity that recognizes its polar headgroup. Functional assays and molecular dynamics simulations delineate the lipid-binding and translocation pathway, revealing how key residues anchor the phosphate of S1P during transport. We also demonstrate that MFSD2B functions as a uniporter and identify residues that mediate the energy-coupling mechanism, uncovering the molecular basis of S1P release from hematopoietic cells.

## Results

### Structure determination of human MFSD2B

MFSD2B is a relatively small membrane protein (∼54 kDa), with most of its mass embedded within the membrane bilayer, making it a challenging target for cryo-EM analysis. To facilitate structural studies of MFSD2B, we adopted a fusion strategy that had previously enabled high-resolution structure determination of other difficult MFS transporters^28,36–38^. Specifically, a maltose-binding protein (MBP) was fused to the N-terminus from the amino acid at position 41 and an MBP-specific binder (DARPin) to the C-terminus of MFSD2B (MFSD2B^EM^ construct). Immunofluorescence analysis showed that MFSD2B^EM^ is predominantly localized to the plasma membrane of human embryonic kidney (HEK293) cells, similar to the wild-type MFSD2B (MFSD2B^WT^) (Extended Data Fig. 1A, B). To ensure that MFSD2B^EM^ remains capable of transporting S1P, we performed a cell-based export assay using cells coexpressing human SphK2 and MFSD2B. The cells were supplied with [³H]-sphingosine, which was subsequently converted to [³H]-S1P by SphK2 and then exported to the medium by MFSD2B. Although the activity was slightly lower than that of MFSD2B^WT^, it is clear that MFSD2B^EM^ retained efficiency to export S1P (Fig. 1A). Consistently, analysis of the [^3^H]-S1P levels in cell pellets indicated that expression of either MFSD2B^WT^ or MFSD2B^EM^ reduced intracellular S1P levels, further supporting the export of S1P from the cells. When cells were supplied with varying concentrations of sphingosine (1–10 μM) for S1P synthesis, we observed dose-dependent S1P export activity (Fig. 1B) and extracellular S1P levels increased progressively over time in both MFSD2B^WT^ and MFSD2B^EM^ (Fig. 1C). Together, these results indicate that the construct modifications have minimal impact on transport function in cells, and that MFSD2B^EM^ largely recapitulates the activity of the wild-type transporter.

**Figure 1:**
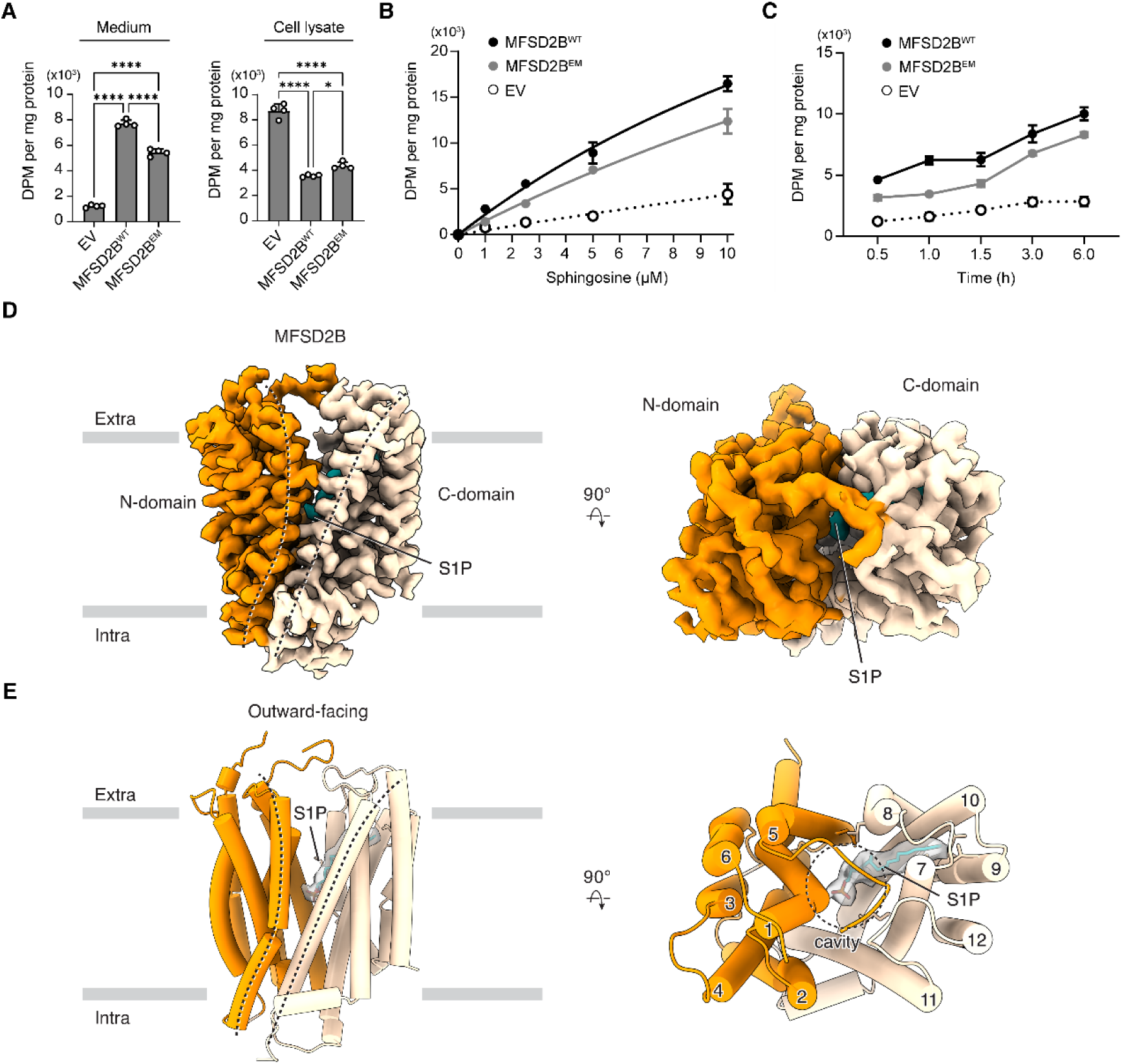
Functional characterization and overall structure of human MFSD2B. (A) Measurement of S1P transport by MFSD2B^WT^ and MFSD2B^EM^ using a cell-based [³H]-S1P export assay. Data are represented as mean ± SEM (n = 4 biological replicates). P-values are calculated using one-way analysis of variance (ANOVA) followed by Dunnett’s multiple comparison test. *^∗∗∗∗^*P < 0.0001; *^∗^*P < 0.05. (B) Dose–response curves for S1P transport activity using different concentrations of sphingosine as the substrate. Data are represented as mean ± SEM (n = 3 biological replicates). (C) Time courses of S1P transport activity using 2.5 μM sphingosine as the substrate. Data are represented as mean ± SEM (n = 3 biological replicates). (D) Cryo-EM density map of MFSD2B in the S1P-bound, outward-facing state. (E) Structural model of MFSD2B. Density corresponding to bound S1P shown in gray where the fatty tail of S1P is inserted into C-domain and the headgroup is pointing to the transport cavity of the transporter.

We determined the structure of MFSD2B^EM^ at 3 Å resolution (Fig. 1D and Extended Data Fig. 2 and Table 1). The quality of the EM density is high, allowing reliable de novo model building for the majority structure of the transporter. MFSD2B adopts a canonical MFS fold comprising twelve transmembrane helices (TMs). TMs 1–6 form the N-domain, and TMs 7–12 form the C-domain (Fig. 1D, E). In this structure, the central cavity between the N- and C-domain was open to the extracellular side, indicating that the transporter is captured in an outward-facing conformation. The protein sample was incubated with exogenous S1P before grid preparation, and a well-defined density corresponding to S1P was observed within the transport cavity of MFSD2B (Fig. 1D, E; gray). In our experiments, n-dodecyl β-D-maltoside (DDM) was used for membrane extraction of MFSD2B before reconstituting into saposin nanoparticles. Detergents used for protein extraction can associate with lipid transporters, as previously reported for SPNS2, where DDM was found to occupy a site within the transporter^31^. Although extensive washing during reconstitution was performed to minimize residual DDM binding to MFSD2B, we sought to exclude the possibility that any remaining detergent accounted for the observed ligand density. Therefore, we extracted MFSD2B using digitonin instead of DDM, and were able to obtain MFSD2B structure at a comparable resolution (3.1 Å and Extended Data Table 1). A similar elongated density was observed in MFSD2B transport cavity (Extended Data Fig. 2F and Table 1). The shape and size of this density are incompatible with a digitonin molecule, indicating that it corresponds to the added S1P.

### S1P binding site and translocation pathway

In our structure, we found that the lipid tail of S1P was inserted deeply into the transporter, binding primarily within the C-domain, with its headgroup pointing toward the central cavity (Fig. 2A, B). In comparison with the previously reported structures of lysolipid transporters in the outward-facing conformation, the lysolipid substrate is either not observed such as in SPNS2 structure^31^ or it is located at the periphery of the transporter such as in MFSD2A and SPNS1 structures^29,35^. Thus, the current structure of MFSD2B with S1P position represents a previously unresolved state of lysolipid translocation.

**Figure 2:**
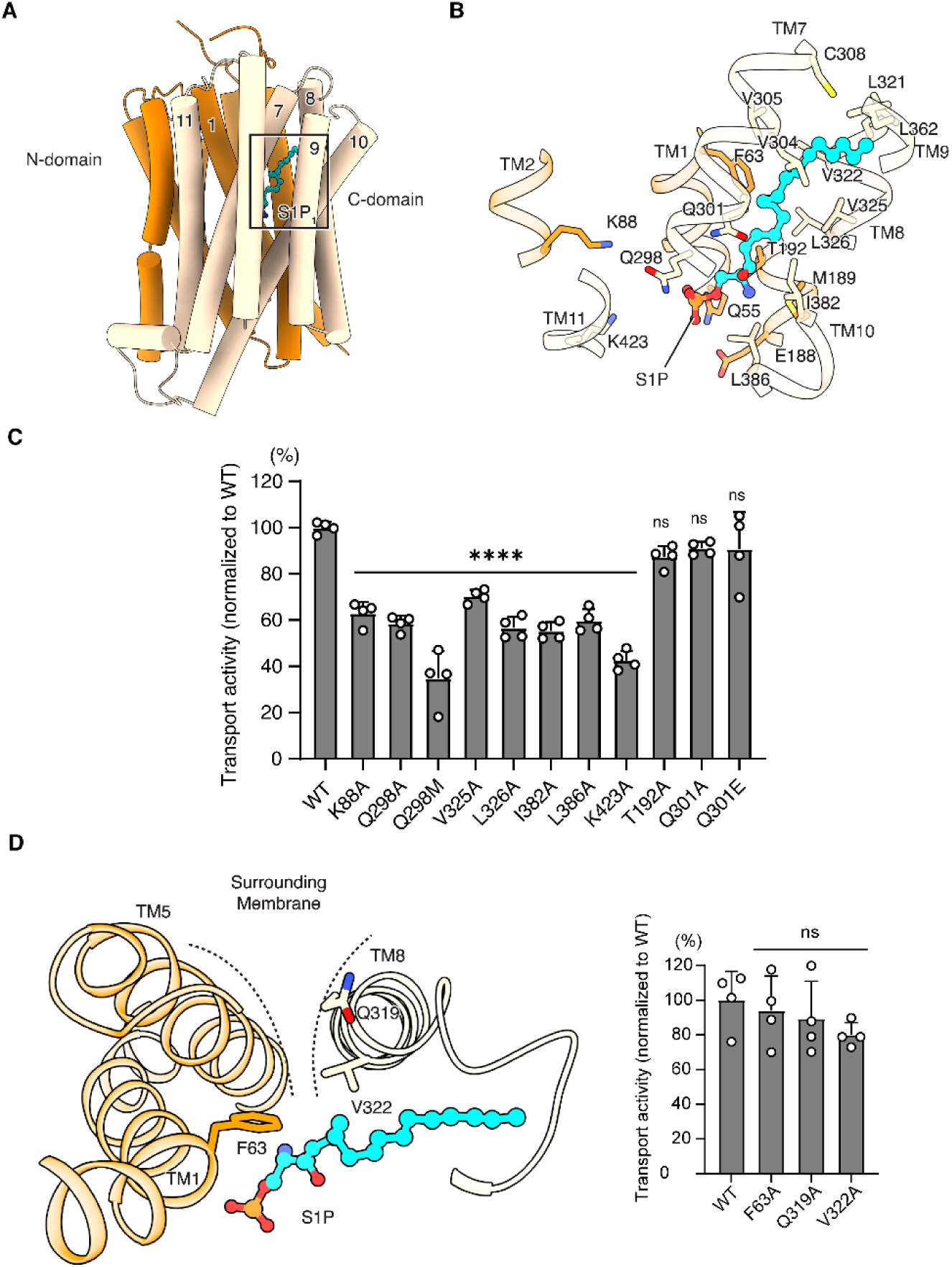
Characterization of S1P binding site in MFSD2B. (A) S1P binding pocket. Green is the fatty tail of S1P, while the headgroup is pointing to the center of the transporter. (B) Detailed interactions between MFSD2B and S1P. Shown are residues that interact with the fatty tail and the headgroup of S1P. (C) [³H]-S1P export activity of MFSD2B variants. The activity is shown as a percentage of the wild type. (mean ± SEM, n = 4 biological replicates). P-values are calculated using one-way ANOVA followed by Dunnett’s multiple comparison test. *^∗∗∗∗^*P < 0.0001; ns, not significant. (D) Left, TM5/TM8 lateral opening in MFSD2B (top view). Right, transport activity of MFSD2B variants mutated in positions predicted to form the gate on TM5 and TM8. The [³H]-S1P export activity is shown as a percentage of the wild type. (mean ± SEM, n = 4 biological replicates). P-values are calculated from one-way ANOVA followed by Dunnett’s multiple comparison test. ns, not significant.

The polar region of S1P is surrounded by a cluster of hydrophilic residues (K88, T192, Q298, Q301, and K423). K423 of MFSD2B is conserved in MFSD2A and has been reported to be critical for binding to the phosphate group of LPC in MFSD2A (Extended Data Fig. 3 and Table 2)^33,35^. To investigate the roles of these residues, we generated a series of mutants that are predicted to disrupt the interaction with S1P headgroup. Protein expression of the variants was largely unaffected (Extended Data Fig. 1C). Interestingly, mutations at K88, Q298, and K423 markedly decreased MFSD2B transport activity (Fig. 2C), indicating that these residues are critical for S1P transport by binding to the headgroup.

The sphingoid backbone of S1P resided in a hydrophobic pocket formed by TM7, TM8, and TM10, which was aligned with hydrophobic conserved residues such as V325, L326, I382, and L386 (Extended Data Fig. 3). To assess the contribution of this region to fatty-tail recognition, we substituted these residues with alanine. These mutations significantly reduced transport activity without affecting protein expression (Fig. 2C and Extended Data Fig. 1C), underscoring the essential role of this hydrophobic pocket in S1P binding and accommodation.

In the extracellular half of MFSD2B, TM5/TM8 and TM2/TM11 form two lateral openings. Similar openings are also present in other lysolipid transporters and commonly serve as lipid translocation pathways. In MFSD2A, lysolipids may enter through either of these openings^26,27,35^, whereas in SPNS1, the TM5/TM8 pathway appears to be preferred and in SPNS2, the TM2/TM11 pathway is favored^28–30^. Because in MFSD2B structure, S1P is located closer to TM5/TM8 opening, we first examined the contribution of this region to its transport activity. Nevertheless, mutations of F63 on TM1, which sits directly in front of the TM5/TM8 opening, and of Q319 and V322 on TM8 had negligible effects (Fig. 2D), suggesting that the TM5/TM8 opening may not be the translocation path for MFSD2B substrate.

To further investigate S1P translocation, we conducted molecular dynamics (MD) simulations. The protein was embedded in a phosphatidylcholine/cholesterol bilayer, with the substrate S1P placed at the position that was observed in the cryo-EM structure. By analyzing the conformational changes of S1P at different time points, we found that the fatty tail stably inserted into the TM7/TM8/TM10 pocket, while its headgroup rotated from the central cavity to the lumen (Fig. 3A–C, Supplementary movie 1). During this process, the two basic residues K88 and K423 also changed their positions to facilitate the interactions with the headgroup of S1P (Fig. 3D). By the end of the simulation, K88 formed a salt bridge with the phosphate group (Fig. 3D), and S1P began to orientate toward the TM2/TM11 opening. Concomitantly, TM2 moved away from TM11, enlarging the lateral opening. The hydrophobic residue F85 on TM2 moved away, possibly creating space to facilitate S1P translocation through the TM2/TM11 gate (Fig. 3E). Importantly, we validated these observations by mutating F85 on TM2 and V419 and F420 residues both on TM11 to alanine. These changes substantially reduced S1P transport (Fig. 3F and Extended Data Fig. 1C). These results indicate that the TM2/TM11 opening in MFSD2B is functionally more relevant for S1P export.

**Figure 3:**
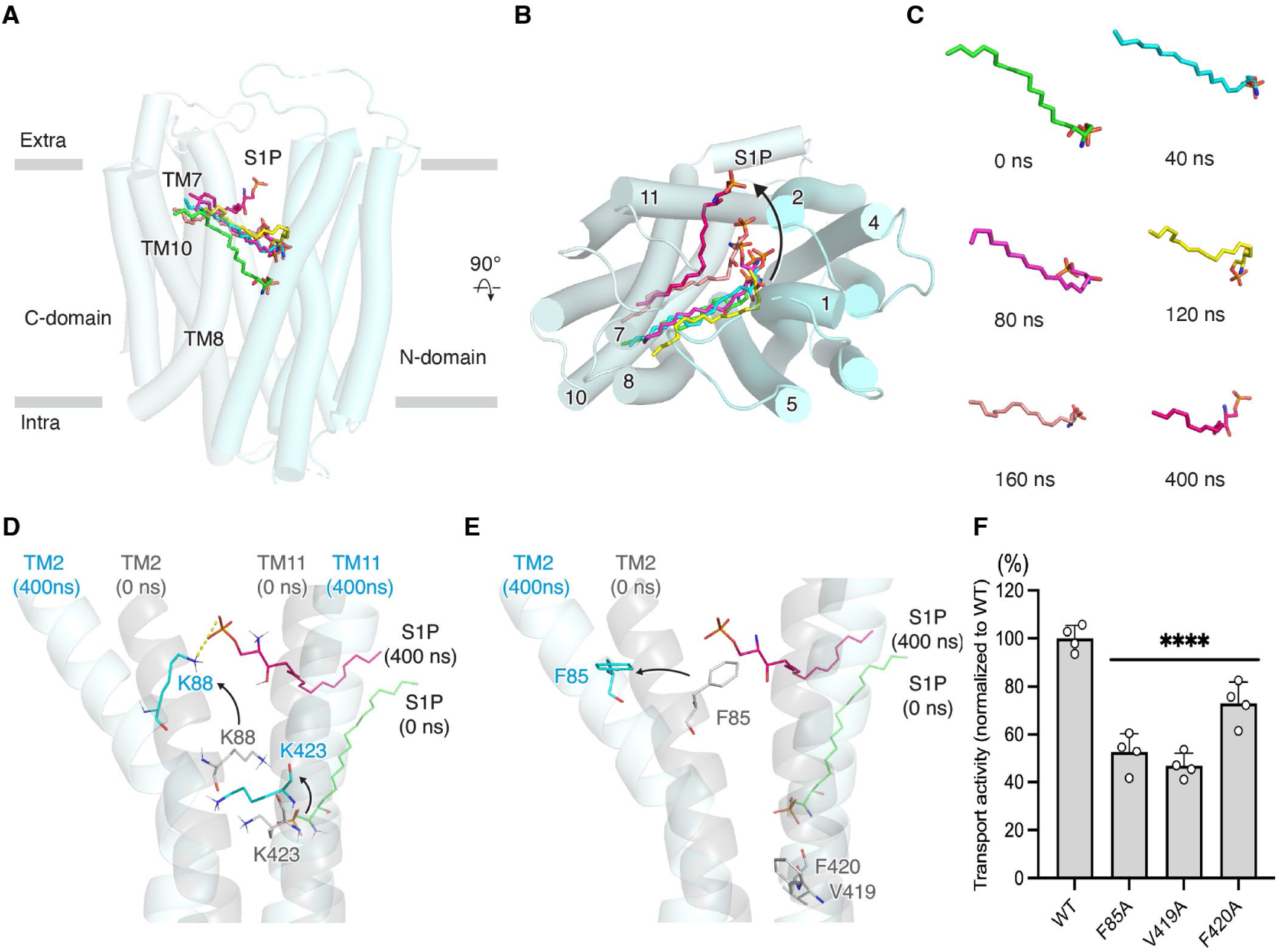
Molecular dynamics simulations of MFSD2B during S1P exit. (**A-B**) Side and top views showing the orientations of S1P in MFSD2B during the 0–400 ns simulations. The headgroup of S1P rotates from the original position (green) to the extracellular side (dark-red). S1P is shown as sticks and colored according to different snapshots. The N-domain and C-domain are colored aquamarine and pale cyan, respectively. (C) Conformations and positions of S1P at various time points during the simulation. S1P is shown as sticks. (D) Comparison of S1P-bound MFSD2B at 0 ns (gray) and 400 ns (cyan). The TM2/TM11 gate opens during MD simulations. Accordingly, the side chains of K88 and K423 reposition to accommodate S1P phosphate group. S1P, K88, and K423 are shown as sticks. (E) Comparison of S1P-bound MFSD2B at 0 ns (gray) and 400 ns (cyan). Residues F85, V419, and F420 move their side chains during S1P exit through the TM2/TM11 gate. S1P, F85, V419, and F420 are shown as sticks. (F) [³H]-S1P export activity of MFSD2B variants mutated in positions predicted to interact with the fatty tail of S1P on TM2/TM11. The activity is shown as a percentage of the wild type. (mean ± SEM, n = 4 biological replicates). P-values were calculated by one-way ANOVA followed by Dunnett’s multiple comparison test. ****P < 0.0001.

### Potential conformational transitions during transport cycle

To gain insights into the structural transitions that occur during the transport cycle of MFSD2B, we next compared our outward-facing structure with its paralog, MFSD2A, which has been captured in outward-facing, outward-occluded, and inward-facing conformations^26,27,32,35^. Relative to the outward-facing conformation of MFSD2A, MFSD2B adopts a more closed configuration (Fig. 4A, top). This is primarily driven by a rotation of the C-domain toward the N-domain on the extracellular side (Fig. 4A, bottom). Individual domain alignments revealed minimal internal rearrangements (Extended Data Fig. 4A, RMSDs of 1.2 Å and 1.0 Å for the N- and C-domains, respectively), consistent with a predominantly rigid-body motion between the two domains. Consequently, transmembrane helices TM8 and TM11 shift closer to TM5 and TM2, respectively, narrowing the extracellular lateral cleft (Extended Data Fig. 4B). Intriguingly, the distance between TM1 and TM7 remains comparable (11.2 Å in both; Fig. 4A, bottom), suggesting a similar degree of opening at this vestibule.

**Figure 4:**
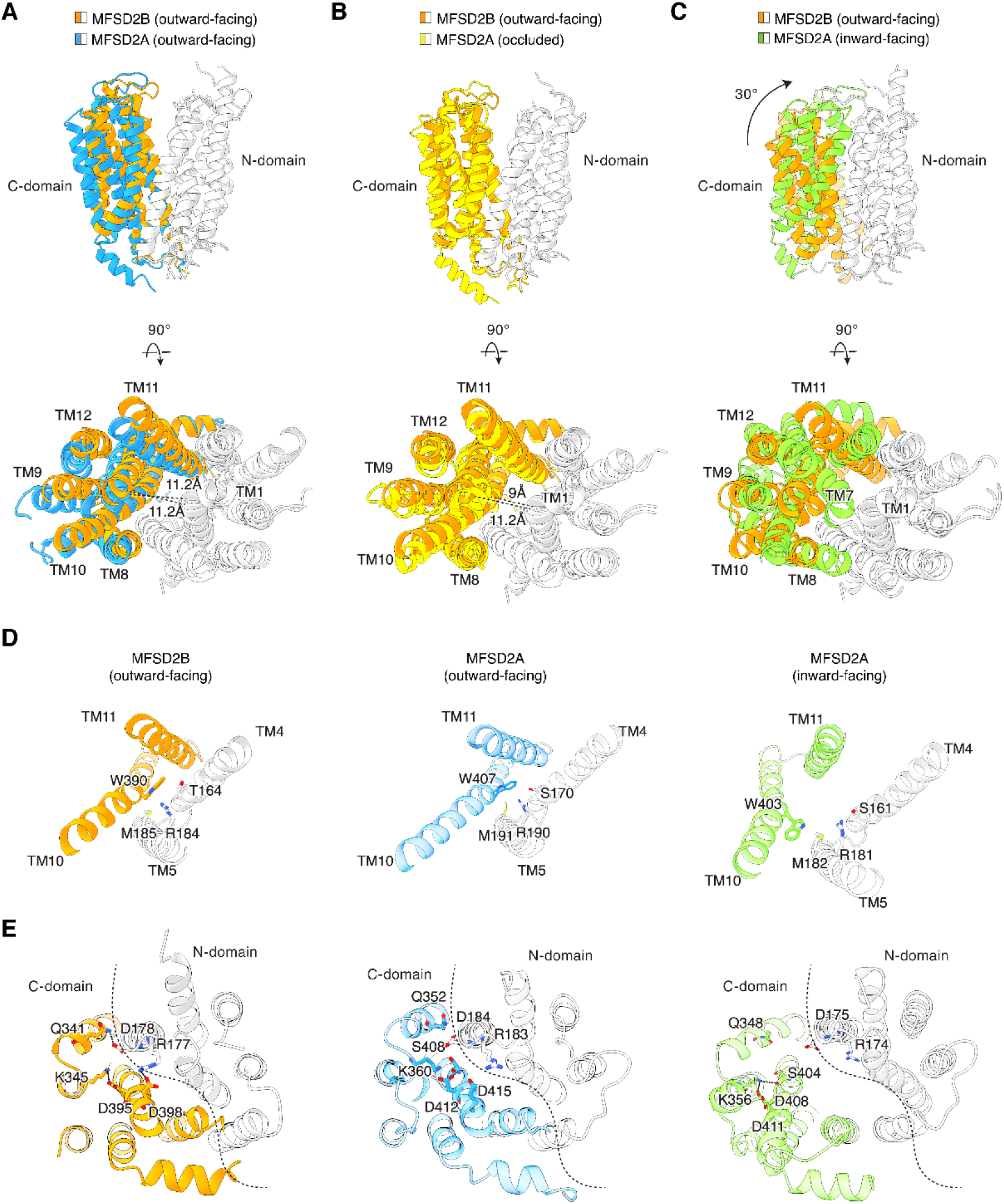
Structural comparison between MFSD2B and MFSD2A. (**A**–**C**) Superimposition of MFSD2B onto MFSD2A structures. The C-domain of MFSD2B is colored orange; the C-domain of MFSD2A is colored blue, yellow, or green. The N-domains of both proteins are shown in gray. PDB codes for MFSD2A: left, 7N98; middle, 7OIX; right, 7MJS. (D) Comparison of the cytoplasmic gates of MFSD2B and MFSD2A, showing conserved residues important for gating. (E) Comparison of inter-domain interactions, viewed from the intracellular side of MFSD2B, showing residues predicted to form inter-domain salt-bridges to stabilize the transporter.

In MFSD2A, TM7 moves closer to TM1 during the transition from the outward-facing to the occluded state (11.2 Å to 9.0 Å; Fig. 4B). It is likely that MFSD2B also undergoes a similar movement, creating a narrow gate that restricts solvent access to the central cavity from the extracellular side (Fig. 4B). To further elucidate how MFSD2B may transition into the inward-facing conformation, we superimposed the outward-facing MFSD2B structure onto the inward-facing MFSD2A structure. This analysis revealed a pronounced rocker-switch-like motion of approximately 30° between the N- and C-domains (Fig. 4C, top). This motion induced reciprocal rearrangements of the transporter at the membrane interfaces. On the extracellular side, TM1 and TM7 constrict the vestibule, thereby closing the extracellular gate (Fig. 4C, bottom), while interactions between TM2–TM11 and TM5–TM8 strengthen to seal the lateral clefts that are thought to mediate lysolipid entry.

This rearrangement propagates through the protein to the intracellular interface, where it repositions key gating residues. A conserved tryptophan (W407 in MFSD2A; W390 in MFSD2B) in TM10 has been shown to act as a pivotal intracellular gate residue in MFSD2A (Fig. 4D, middle). In the outward-facing conformation, it engages S170 residue (T164 in MFSD2B) on TM4 and M191 (M185 in MFSD2B) and R192 (R184 in MFSD2B) residues on TM5, occluding access to the cytosol. The same interaction of these conserved residues in MFSD2B is also observed (Fig. 4D, left). However, this interaction is disrupted in the inward-facing conformation (Fig. 4D, right), permitting gate opening.

The structural transition is accompanied by a remodeling of electrostatic contacts. Several inter-domain interactions, such as the R177–D398 salt bridge in MFSD2B and the R183–D415 salt bridge in MFSD2A, stabilize the outward-facing state and contribute to the closure of the intracellular vestibule (Fig. 4E, left and middle), whereas the outward-to-inward transition breaks these interactions to permit cytoplasmic opening (Fig. 4E, right). Notably, the MFSD2B structural model predicted by AlphaFold2 adopts an inward-facing conformation. We therefore compared our structure with the AlphaFold2 model^39^ (Extended Data Fig. 4C). The conformational transition appeared similar to that has been observed when comparing MFSD2B with MFSD2A in different states: a rocker-switch like motion involving the opening of the intracellular gate, exemplified by the movement of W390 residue (Extended Data Fig. 4D), and the disruption of inter-domain hydrophilic interactions (Extended Data Fig. 4E). Although determining the structure of MFSD2B in additional conformational states will be essential to fully delineate its transport cycle, the present analysis provides a plausible framework for understanding its conformational transitions.

### MFSD2B is an S1P uniporter

Unlike MFSD2A, which uses sodium to drive LPC/LPE import, MFSD2B transport function is independent of sodium^6^. Its activity, however, appears to be influenced by extracellular pH^6^. To examine whether MFSD2B-mediated S1P transport is dependent on a proton gradient, we measured [³H]-S1P release under different pH conditions. MFSD2B was capable of exporting S1P across a broad pH range, although its activity was slightly reduced at pH 5.5 (Fig. 5A). These results suggest that while MFSD2B activity may be affected by extracellular pH, S1P transport by MFSD2B does not require a transmembrane proton gradient. The independence from sodium and proton in MFSD2B is reminiscent of the transport properties of SPNS2, which functions as a uniporter^28^. If MFSD2B also operates as a uniporter, it should mediate S1P influx when the extracellular concentration exceeds that inside the cell. Indeed, under such experimental conditions, we observed S1P uptake mediated by MFSD2B (Fig. 5B), consistent with a previous study^40^. We also reconstituted purified MFSD2B into liposomes and performed S1P uptake assay under conditions lacking any ion gradient across the membrane. Our results showed that MFSD2B remained active and was able to import [³H]-S1P and NBD-S1P (Fig. 5C, D and Extended Data Fig. 5A–C). These results provide strong evidence that MFSD2B functions as a uniporter that facilitates S1P transport driven by the concentration gradient.

**Figure 5:**
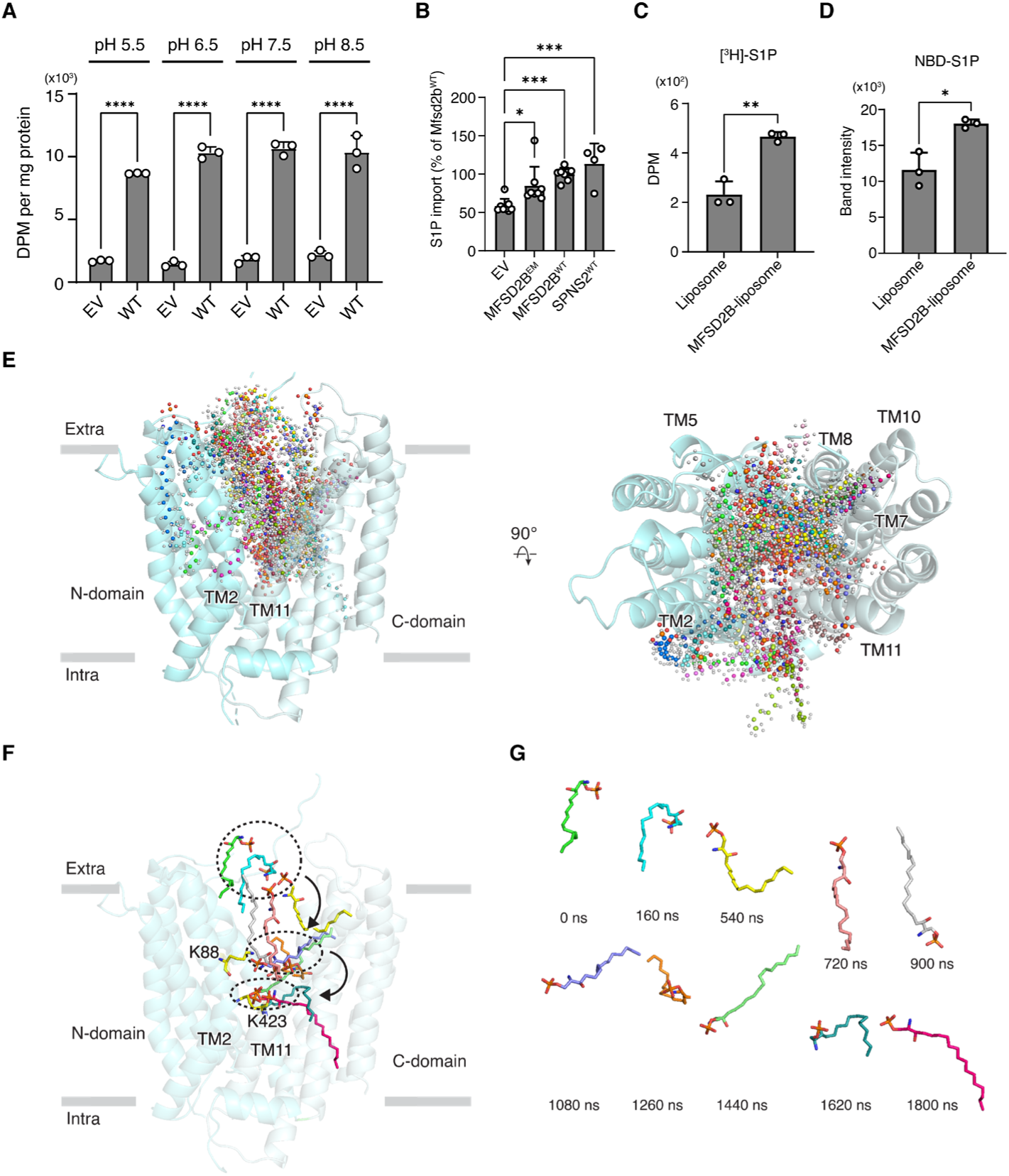
MFSD2B functions as a uniporter. (A) Effect of extracellular pH on [³H]-S1P export activity. Data are presented as mean ± SEM (n = 3 biological replicates). P-values were calculated using one-way ANOVA followed by Dunnett’s multiple-comparison test. ****P < 0.0001. (B) MFSD2B import activity of S1P(d18:1) measured by LC–MS/MS analysis and shown as percentage relative to the wild-type MFSD2B (mean ± SEM, n = 4–8 biological replicates). SPNS2, a S1P uniporter, was used as a positive control. P-values were calculated using one-way ANOVA followed by Dunnett’s multiple-comparison test. ***P < 0.001; *P < 0.05. **(C**, **D)** Proteoliposome-based transport assays. Controls and MFSD2B-containing liposomes were pre-loaded with 0.1% BSA. [³H]-S1P (C) or NBD-S1P (D) were used as the substrate. [³H]-S1P signal from the proteoliposomes was detected using scintillation counter, while NBD-S1P signal was analyzed by TLC and the fluorescent band was quantified. Data are presented as mean ± SD (n = 3 biological replicates). P-values were determined by two-tailed t-test. **P < 0.01; *P < 0.05. (E) Side and top views showing movements of S1P atoms (shown as spheres) from MD trajectory frames. The N-domain and C-domain of MFSD2B are colored aquamarine and pale cyan, respectively. (F) Positions of S1P at selected time points in the S1P-bound MFSD2B during simulations where S1P was placed at the extracellular side (green). The high tendency of S1P movements accessing to TM2/TM11 gate and the engagement of K88 and K423 residues were observed. K88 and K423 are shown as sticks and colored yellow. (G) Conformations of S1P at various time points from extracellular side (green) to the position where the phosphate group is bound to K423 and the fatty tail is inserted deep into C-domain.

Since MFSD2B is a uniporter capable of importing S1P when extracellular concentrations are high, we also performed MD simulations where S1P was placed on the extracellular side of MFSD2B to investigate how it enters the transporter (Supplementary movie 2). Umbrella sampling with 60 windows yielded a cumulative sampling time of 1.8 μs. Superimposing snapshots of S1P along the trajectory revealed that while its fatty tail explored both the TM5/TM8 and TM2/TM11 lateral openings, the tendency insertion of the substrate into the TM2/TM11 pathway occurred more frequently (Fig. 5E). During the early stage (0–540 ns) of translocation, S1P primarily engaged the TM2/TM11 cleft, suggesting this route serves as the initial entry point (Extended Data Fig. 6A, B). Once the headgroup of S1P bound to the central cavity, the fatty tail remained in the hydrophobic pocket formed by TM7, TM8, and TM10 (Extended Data Fig. 6C, D), consistent with our structural and mutagenesis data (Fig. 2B, C). Together, our functional and simulation data indicate that the TM2/TM11 lateral opening may serve as the primary route for lipid entry and release in MFSD2B.

In these MD simulations, we observed a stepwise movement of S1P, in which its phosphate group appeared to engage the charged residues K88 and K423 as sequential anchors to move deeper into the cavity (Fig. 5F, G). A similar mechanism has also been reported for MFSD2A and SPNS2^30,33^. In the first 0–540 ns, the headgroup of S1P predominantly remained near the extracellular side of the transporter, while its fatty tail exhibited a waving motion. Between 720 and 1260 ns, the headgroup moved toward the central cavity, strengthening its interaction with K88. Subsequently, between 1440 and 1800 ns, the headgroup of S1P descended to move deeper into the cavity, bringing the phosphate group into proximity with K423 (Fig. 5F, G). To elucidate the roles of key residues in the transport process, we performed potential of mean force (PMF) calculations to characterize the free energy landscape of S1P movement ^41^. Four representative conformations at key turning points (510, 900, 1080, and 3960 ns) revealed that residues K88 and K423 around the central cavity play crucial roles in recognizing S1P by forming salt bridges with phosphate group in S1P (Extended Data Fig. 7). Additionally, polar residues Q55, Q160, and E188 contribute to S1P recognition through hydrogen-bonding interactions. These interactions stabilize S1P within the central cavity and promote its progressive translocation in the transporter.

### Coupling mechanism in MFSD2B

Previous studies have shown that two aspartic acid residues of mouse MFSD2A (D92 and D96) are critical for Na⁺ binding^35^. In MFSD2B, these positions correspond to G91 and D95 (Extended Data Fig. 3). This divergence, where the first aspartate becomes a glycine in MFSD2B, likely reflects an evolutionary adaptation, allowing the transporter to operate efficiently without relying on sodium or proton gradients to export S1P in hematopoietic cells. To test this hypothesis, we generated the G91D mutant and examined its activity in buffers at pH 5.5 and pH 7.5, with or without sodium (NaCl or choline chloride, respectively). In sharp contrast to MFSD2B^WT^, this mutant was active at pH 5.5 but inactive at pH 7.5 regardless of sodium or choline chloride presence (Fig. 6A and Extended Data Fig. 1C). Thus, although this single substitution does not confer sodium dependence, it remarkably converts MFSD2B into a proton-dependent transporter that possibly replies on proton influx to drive S1P export. To evaluate the role of these aspartic acids in proton dependence, we tested the G91D/D95A which has a single aspartate like MFSD2B^WT^. The results showed that the mutant remained active, especially at normal pH (Fig. 6B and Extended Data Fig. 1C). D95A, which lacks both acidic residues, was inactive in the same condition. Thus, the mutation of D to G at position 91 in MFSD2b likely abolishes or reduces the cation-dependent function of the transporter. Only when both aspartic residues are present, as in MFSD2B^G91D^, does the transporter function like a proton/S1P antiporter.

**Figure 6:**
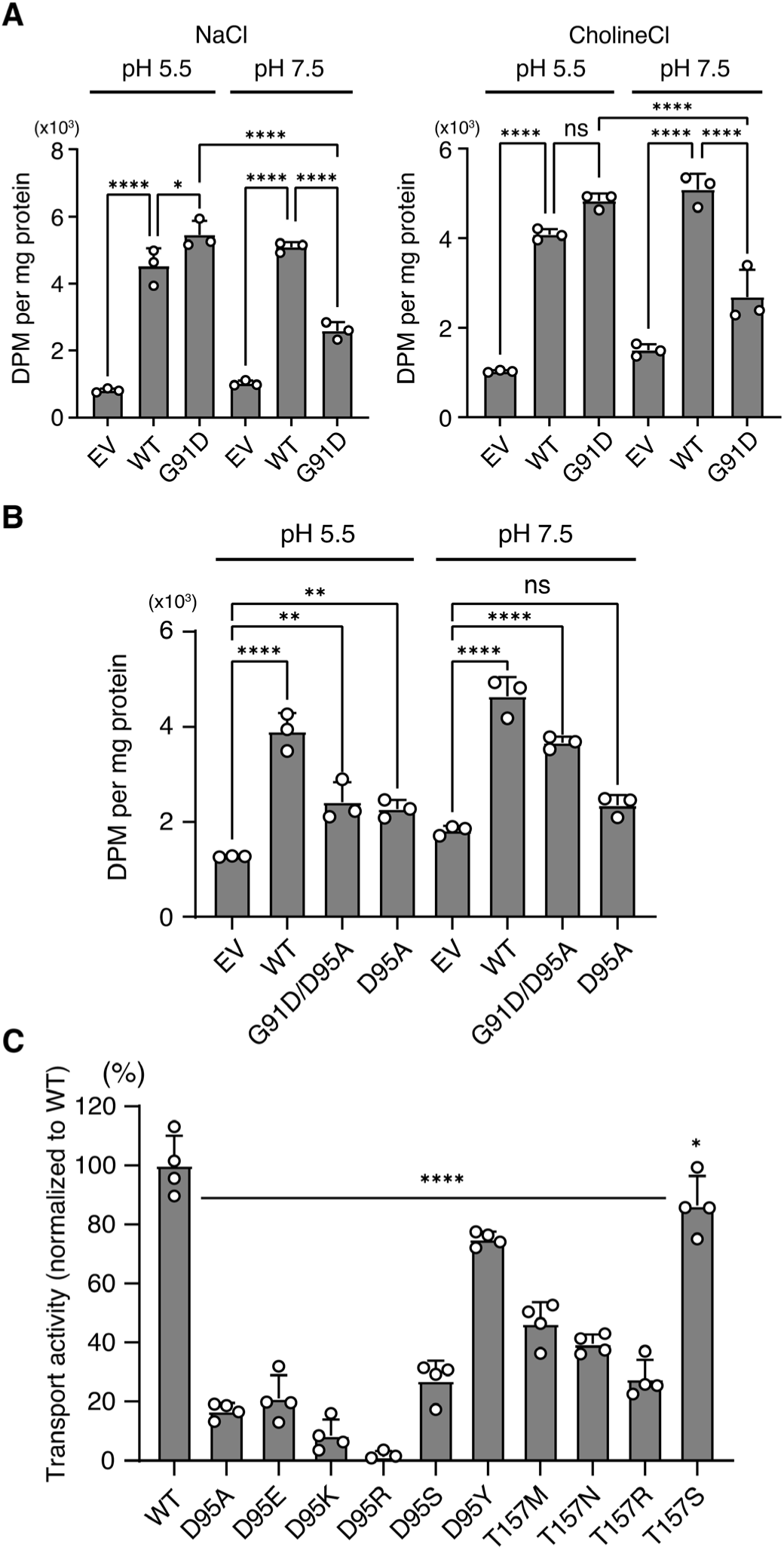
Determination of MFSD2B coupling machinery. (A) Cellular [³H]-S1P export from MFSD2B^WT^ and indicated mutants in pH 5.5 or pH 7.5 buffer supplemented with NaCl or choline chloride. Data are presented as mean ± SEM (n = 3 biological replicates). P-values were calculated using one-way ANOVA followed by Dunnett’s multiple-comparison test. ****P < 0.0001; *P < 0.05; ns, not significant. (B) Cellular [³H]-S1P export from MFSD2B^WT^ and indicated mutants in pH 5.5 or pH 7.5. Data are presented as mean ± SEM (n = 3 biological replicates). P-values were calculated using one-way ANOVA followed by Dunnett’s multiple-comparison test. ****P < 0.0001; **P < 0.01; ns, not significant. (C) Cellular [³H]-S1P export from MFSD2B^WT^ and indicated mutants. The activity is shown as percentages relative to the wild type. Data are presented as mean ± SEM (n = 4 biological replicates). P-values were calculated using one-way ANOVA followed by Dunnett’s multiple-comparison test. ****P < 0.0001. EV, empty vector–transfected cells (co-transfected with SphK2).

In MFSD2A, T172 also participates in the Na⁺-binding network^27^. The corresponding residue in MFSD2B is T157 (Extended Data Fig. 3). We therefore performed additional mutagenesis to examine the importance of side chain of D95 and T157. Substitution of either residue with different amino acids markedly reduced S1P export, with the exception of D95Y and T157S, which remained partially active (Fig. 6C and Extended Data Fig. 1C). These results suggest that both the side chain length and polar properties of D95 and T157 residues are required for MFSD2B function. Because D95Y preserved substantial activity, suggesting that the hydroxyl group of tyrosine can replace the carboxylic group of aspartic acid. Thus, we tested whether MFSD2B^G91D/D95Y^ could function as a proton/S1P antiporter, similar to MFSD2B^G91D^ mutant. However, this mutant behaved similarly to the wild-type protein and was not inactive at high extracellular pH (Extended Data Fig. 8A). Thus, it appears that two aspartic residues are required for cation dependence in this family of transporters.

To understand how the G91D substitution enables proton-dependent S1P transport, we used AlphaFold2 to generate a structural model of MFSD2B^G91D^ and predict the potential interactions of D91 with nearby residues. We found that the amine group of K423 formed two salt bridges with the carboxylic acids of D91 and D95, respectively (Extended Data Fig. 8B). These salt bridges are absent in the wild-type transporter, where the amine group of K423 instead points toward the central cavity (Extended Data Fig. 8B). Since K423 is a critical residue for S1P binding (Figs. 2C, 3D, and 5F), we speculated that the loss of activity in G91D at pH 7.5 arises from the forming of these salt bridges that restrict K423 from engaging S1P. At low pH, protonation of D91 and D95 disrupts these interactions, allowing K423 to bind S1P, thus enabling the transport activity. Collectively, these findings identify the D91–K423–D95 triad as a key structural element that enables proton coordination and S1P transport in the G91D mutant. The G91D substitution unveils how subtle rearrangements in local charge networks can reprogram the transport mechanism of MFSD2B.

## Discussion

Here we provide a structural and mechanistic basis for MFSD2B-mediated S1P transport. By combining cryo-EM, functional assays, and molecular dynamics simulations, we reveal how this transporter accommodates and translocates S1P. Our structural data revealed MFSD2B in an outward-facing conformation. We were unable to capture it in an inward-facing conformation, despite extensive efforts using varying substrates, detergents, lipid compositions, and nanodisc scaffolds. Nevertheless, our MD simulations and comparison with its closest paralog MFSD2A suggest that MFSD2B also operates via a rocker-switch mechanism (Figs. 3, 4).

Our data show that TM7/TM8/TM10 hydrophobic pocket forms a cleft with a crucial role in S1P fatty tail binding. Previous studies on MFSD2A suggest both TM5/TM8 and TM2/TM11 openings can potentially serve as gates for its lipophilic ligand’s binding and entry. In contrast, in our MD simulations of MFSD2B, S1P shows a higher propensity towards the TM2/TM11 gate, which widens substantially for S1P exit. Mutagenesis results of residues on TM2/TM11 but not TM5/TM8 strongly affect the transport activity. This suggests that S1P might exit through the gate between TM2 and TM11 in MFSD2B.

Two positively charged residues, K88 and K423, line the central cavity and engage the phosphate group of S1P in a stepwise manner during translocation, akin to mechanisms observed in MFSD2A and SPNS2. The binding of S1P is sufficient to trigger conformational transitions in MFSD2B, enabling lipid translocation across the membrane. In order to import LPC/LPE, human MFSD2A requires co-transport of sodium, which binds to residues D92 and D97 residues. In contrast, MFSD2B^WT^ does not rely on proton nor Na^+^ gradient to transport S1P and residue G91 (the position equivalent to MFSD2A D92) is likely responsible for this difference between the two related transporters. Indeed, mutating G91 to D converted MFSD2B into a proton-dependent transporter, with G91D and D95 (equivalent to MFSD2A D97) playing crucial roles in the proton-sensing mechanism, as they would coordinate with K423 to form salt bridges thus interfering with S1P binding at physiological pH. At lower pHs, D91 and D95 would be protonated, disrupting salt bridges and releasing K423 to engage with the phosphate group of S1P. Thus, the appearance of G91 in MFSD2B has allowed it to adopt a facilitated diffusion mechanism to transport S1P.

In summary, our work reveals the molecular basis by which MFSD2B operates to transport S1P in hematopoietic cells. Further work will be required to fully understand how S1P flips its orientation within the transporter during this transport cycle and how it is delivered to albumin or HDL, which are the major carriers of S1P in blood.

## Methods

### S1P release assay in HEK293 cells

HEK293 cells (ATCC) were cultured in Dulbecco’s Modified Eagle Medium (DMEM) (Gibco, 11995-065) supplemented with 10% Fetal Bovine Serum (FBS) and 1% penicillin-streptomycin (Gibco) at 37 °C in 5% CO_2_. Cells at 80% confluency were transfected with the indicated plasmids using Lipofectamine 2000 Transfection Reagent in accordance with the manufacturer’s manual (Invitrogen, 11680-019).

HEK293 cells were co-transfected with plasmid carrying human Sphk2 gene cloned in pcDNA3.1 vector and empty vector (pcDNA3.1 or pIRES2), human MFSD2B^WT^, MFSD2B^EM^ or mutant constructs. After 24 hours of transfection, the cells were incubated in plain DMEM containing 2.5 μM [^3^H]-sphingosine with a cocktail of phosphatase inhibitors (5 mM sodium fluoride, 1 mM semicarbazide and 10 mM sodium glycerophosphate) for three hours. Subsequently, the medium was changed to plain DMEM supplemented with 0.5% BSA and the phosphatase inhibitors to facilitate the release of [^3^H]-S1P for additional 2-hours. Then, medium was harvested for S1P isolation process and quantification of radioactive signals by a Liquid Scintillation Counter (LSC Tri-carb 5110R, Perkin Elmer, USA). The remaining medium of each group was removed, the cells were washed with plain DMEM and then lyzed with 400 μL RIPA and shaken for 30 min for cell lysate signal quantification or protein normalization.

For the dose curve assay, the transfected cells were incubated with indicated concentrations of [^3^H]-sphingosine. For the time course assay, the supernatant after changing the 0.5% BSA containing medium was collected at indicated time. For testing the effect of different pHs on the MFSD2B transport activity, these following transport buffers were used: pH 5.5 (140 mM NaCl, 20 mM MES, 2 mM CaCl_2_, 1 g/L D-glucose), pH 6.5 (140 mM NaCl, 20 mM MES, 2 mM CaCl_2_, 1 g/L D-glucose), pH 7.5 (140 mM NaCl, 20 mM HEPES, 2 mM CaCl_2_, 1 g/L D-glucose), pH 8.5 (140 mM NaCl, 20 mM Tris, 2 mM CaCl_2_, 1 g/L D-glucose). For testing the effect of Na cation on the MFSD2B transport activity, these following transport buffers were used: NaCl-pH 5.5 (140 mM NaCl, 20 mM MES, 2 mM CaCl_2_, 1 g/L D-glucose), Choline chloride-pH 5.5 (140 mM choline chloride, 20 mM MES, 2 mM CaCl_2_, 1 g/L D-glucose), NaCl-pH 7.5 (140 mM NaCl, 20 mM HEPES, 2 mM CaCl_2_, 1 g/L D-glucose), Choline chloride-pH 7.5 (140 mM choline chloride, 20 mM HEPES, 2 mM CaCl_2_, 1 g/L D-glucose).

All functional experiments were repeated at least twice with 3 biological replicates on different days. Similar results were obtained. Statistical analyzes of data were performed using GraphPad Prism 10. P value below 0.05 was considered as significance. Quantification methods and tools used are described in each relevant section of the methods or figure legends.

### Western blotting

HEK293 cells were seeded onto 12-well plates and transfected with indicated plasmids. After 24 hours incubation, the cells were washed with cold phosphate buffered saline (PBS) and lyzed with RIPA supplemented with protease inhibitor (Roche). Equal amount of total protein was loaded on 10% SDS-PAGE gel. The proteins were then transferred to 0.45 µm nitrocellulose membrane. Subsequently, the membranes were blocked with TBST buffer (Tris-buffered saline with 0.1% Tween 20) containing 5% skim milk at room temperature for 1 hour. The membranes were blotted with primary antibody (in-house generated MFSD2B, dilution 1:500) overnight in cold room, washed with TBST three times. Then the membranes were incubated with IRDye 800CW Goat anti-Rabbit IgG (H + L) secondary antibody (dilution 1:5000) for 1 hour at room temperature, washed with TBST three times, and detected using ChemiDoc (Biorad). The membranes were also re-probed with mouse monoclonal GAPDH antibody as loading control.

### Immunostaining

HEK293 cells were grown on cover glass for overnight and transfected with respective plasmids as described. After 24 hours transfection, cells were washed and fixed in 3.7% methanol-free formaldehyde in PBS for 15 minutes and permeabilized for 3 minutes in PBS with 0.1% Triton-X. After 15 minutes blocking in PBS with 5% Natural Goat Serum (NGS), the cells were incubated with primary antibody against MFSD2B (1:250 diluted in 5% NGS) for 1 hour, followed by incubation in Goat Anti-Rabbit IgG (H+L), Alexa Fluor 555 conjugate Antibody (1:500 diluted in 5% NGS) for another one hour. The cells were counterstained with Hoechst 33342 to visualize the nuclei. The cells were visualized under Zeiss LSM 710 Confocal Microscope at 100x magnification.

### Expression and purification of human MFSD2B^EM^

For MFSD2B^EM^, a maltose-binding protein (MBP) was inserted at the N-terminus, followed by a helical linker, truncated MFSD2B protein (residues 41–504), a DARPin (off7) at the C-terminus^42^, and a mVenus tag. Stable Expi293 cell lines expressing MFSD2B^EM^ were generated using the PiggyBac transposon system. After puromycin selection, cells exhibiting strong fluorescence were enriched by fluorescence-activated cell sorting, yielding a population suitable for high-level MFSD2B expression^43^.

The pellet of cells expressing MFSD2B was resuspended using hypotonic buffer (10 mM NaCl, 1 mM MgCl_2_, 20 mM HEPES pH 7, benzonase, and protease inhibitors) for 30 min. The cell lysate was then spun at 39,800 g for 35 mins to sediment crude membranes. The membrane pellet was mechanically homogenized and solubilized in extraction buffer (20 mM DDM, 4 mM CHS, 300 mM NaCl, 20 mM HEPES pH 7, 1 mM TCEP, and protease inhibitors) for 1.5 hours. For certain experiments, 1.5% digitonin instead of DDM was used for solubilization. Solubilized membranes were clarified by centrifugation at 39,800 g for 40 min. The supernatant was applied to a GFP nanobody-coupled sepharose resin^44^, which was subsequently washed with 10 column volumes of wash buffer A (0.05% digitonin, 150 mM NaCl, 20 mM HEPES pH 7, 4 mM NaATP, 4 mM MgCl_2_, and 1 mM TCEP), followed by 7 column volumes of wash buffer B (0.05% digitonin, 150 mM NaCl, 20 mM HEPES pH 7, and 1 mM TCEP). The washed resin was incubated with 3C protease for 2 hours at a target protein to protease ratio of 40:1 (w/w) to cleave off mVenus and release the protein from the resin. The protein was eluted with wash buffer B, concentrated, and further purified by gel-filtration chromatography using a superose 6 Increase column equilibrated with Buffer B. The peak fractions were concentrated and incubated with saposin, at a molar ratio of 1:15 (MFSD2B: saposin). After 30 mins, γ-cyclodextrin was added to the sample to remove digitonin and initialize the nanoparticle reconstitution. After 14 hours, the reconstituted protein was buffer exchanged in a 100 kDa centrifugal filter to remove the excess saposin and further purified by gel-filtration chromatography using a superose 6 Increase column equilibrated with SEC buffer (150 mM NaCl, 20 mM HEPES pH 7, and 1 mM TCEP). Peak fractions were pooled and incubated with 10 μM S1P for 1 hour, then concentrated for cryo-EM experiments.

### Proteoliposome-based transport assay

Proteoliposomes were prepared as previously reported^30^. Liposomes were generated by combining 80 mol% POPC and 20 mol% cholesterol in buffer A (20 mM HEPES, 150 mM NaCl, pH 7.5). The lipid mixture was hydrated and extruded to form unilamellar vesicles, which were subsequently supplemented with 0.1% (w/v) fatty acid–free bovine serum albumin (BSA). To reconstitute MFSD2B, 50 μg of purified protein was incorporated into 160 μL of preformed liposomes in the presence of 0.11% Triton X-100. The detergent was removed using SM-2 Bio-Beads (Bio-Rad) under gentle agitation at 4 °C for 2 h. Following detergent removal, the resulting proteoliposomes were pelleted and washed twice with buffer A, then resuspended in 320 μL of the same buffer. Liposomes processed in parallel without protein insertion served as negative controls. For uptake assays, MFSD2B proteoliposomes or control liposomes were incubated with 2.5 μM [^3^H]-S1P or 2.5 μM NBD-S1P at 37 °C for 1 h. Reactions were terminated by the addition of 1 mL cold buffer A, followed by three rounds of centrifugation and washing. [^3^H]-S1P uptake was quantified by Liquid Scintillation Counter. For fluorescent assays, NBD-labeled lipids were extracted from liposomes using methanol:chloroform (2:1, v/v), spotted onto silica gel TLC plates, and visualized using a fluorescence imaging system (ChemiDoc MP, Bio-Rad).

### Cryo-EM sample preparation and data acquisition

Protein samples were concentrated to 10 mg/mL, and 0.4 mM fluorinated octyl maltoside and an MBP-specific nanobody (at a molar ratio of 1:1.2, MFSD2B:saposin) were added^45^. The samples were then centrifuged to remove insoluble S1P prior to grid preparation. Aliquots of 3.5 μl protein samples were applied to plasma-cleaned UltrAuFoil R1.2/1.3 300 mesh grids (Quantifoil). The grids were blotted for 1.5 s and plunged into liquid ethane using a Vitrobot Mark IV (FEI) operated at 10 °C and 100% humidity. The grids were loaded onto a 300 kV Titan Krios transmission electron microscope for data collection. Raw movie stacks were recorded with a K3 camera at a physical pixel size of 0.649 Å per pixel and a nominal defocus range of 1.1–2.1 μm. The exposure time for each micrograph was 1.8 s, dose-fractionated into 60 frames with a dose rate of ∼1.0 e^−^/Å^2^/s. The data collection parameters are summarized in Extended Data Table 1.

### Cryo-EM data processing

The image stacks were gain-normalized and corrected for beam-induced motion using cryoSPARC4^46^. Defocus parameters were estimated from motion-corrected micrographs in cryoSPARC4. Micrographs unsuitable for further analysis were excluded after manual inspection. Particle picking (blob, template, and TOPAZ pickers^47^) and two-dimensional classification were performed in cryoSPARC4. Selected particles from both pickers were merged, and duplicates were removed. Iterative three-dimensional classification was then carried out, followed by ab initio reconstruction and heterogeneous refinement to eliminate suboptimal particles. The selected particles were refined using non-uniform refinement^48^, followed by local refinement with soft masks encompassing MFSD2B to further improve map quality. Mask-corrected FSC curves were calculated in cryoSPARC4, and resolutions were reported based on the 0.143 FSC criterion.

### Model building and refinement

An initial model of human MFSD2B was generated by AlphaFold2. This model was docked into the density map using Chimera^49^. The model was then refined iteratively using Coot^50^, ISOLDE^51^, and Phenix^52^. Structural model validation was performed using Phenix and MolProbity^53^. Figures were prepared using ChimeraX^54^.

### Molecular dynamic simulation

All molecular dynamics simulations were performed using the GROMACS 2024.3 software^55^. Our structure was embedded in a lipid bilayer with POPC and cholesterol in a ratio of 3:1 prepared by CHARMM-GUI^56^, and solvated in TIP3P water with 150 mM NaCl. The CHARMM36m force field was used^57^. Long-range electrostatic interactions were calculated by particle-mesh Ewald^58^. Van-der-Waals interactions were cut-off beyond a distance of 1.2 nm. The LINCS algorithm was adopted to constrain the bonds involving hydrogen atoms^59^. During equilibration, a constant temperature of 310.15K was maintained with the velocity-rescale^60^. Constant pressure of 1 bar was established with the semi-isotropic canonical rescale^61^. In the production runs, the same algorithms for controlling the temperature and pressure were used. MD simulations with our structure of MFSD2B in complex with S1P were conducted for a total running time of 400 ns. For simulations where S1P was placed outside the cavity, umbrella sampling with the Gromacs weighted histogram analysis method (WHAM) was applied to evaluate a one-dimensional PMF^62^. Steered molecular dynamics simulations were initiated using the center of mass between the protein and S1P as the collective variable, with a pulling rate of 0.01 nm/ps. Initial configurations were extracted along the Z-axis of the system at intervals of 0.05 nm. Each window was energy-minimized using the steepest descent until converged, then subjected to 10 ns of equilibration in the isothermal isobaric ensemble with umbrella harmonic restraints on. A total of 60 windows of 30 ns in length for a total of 1.8 µs sampling was obtained. The trajectories and conformations were analyzed and visualized in Pymol. Videos were generated by Pymol.

## Data availability

The cryo-EM maps have been deposited in the Electron Microscopy Data Bank (EMDB) under accession codes EMD-74413 (DDM extraction) and EMD-74414 (digitonin extraction). The atomic coordinates have been deposited in the Protein Data Bank (PDB) under accession codes 9ZLV (DDM extraction) and 9ZLW (digitonin extraction).

## Acknowledgements

We thank scientists in the Cryo-EM Center of St. Jude Children’s Research Hospital for their support in data collection. We thank members of the Nguyen lab including Dr. Nguyen. Q.T. and Yang X. for making several mutants. We also thank members of the Lee lab and Dr. Y. Niu for helpful discussions; Dr. I. Chen for editing the manuscript. We thank Dr. Zhao L. for the support with MD simulations. This work is supported by National Institutes of Health (R01GM143282 R01NS133147) and ALSAC to C.-H.L and Singapore Ministry of Education T2EP30221-0012 (T2EP30221-0006), T2EP30123-0014 (T2EP30123-0016), MOE Tier 1, and STDR pre-pilot grants to L.N.N..

## Authors Contributions

S.A. performed biochemical and structural experiments. M.H. performed functional experiments. Y.D. contributed to structural analysis. M.H. and X.Y. performed MD simulations. C.-H.L. and L.N. N. conceived the research and supervised the project. C.-H.L., L.N. N. and M.H. wrote the manuscript with input from all authors.

---

**Extended Data Fig. 1:**
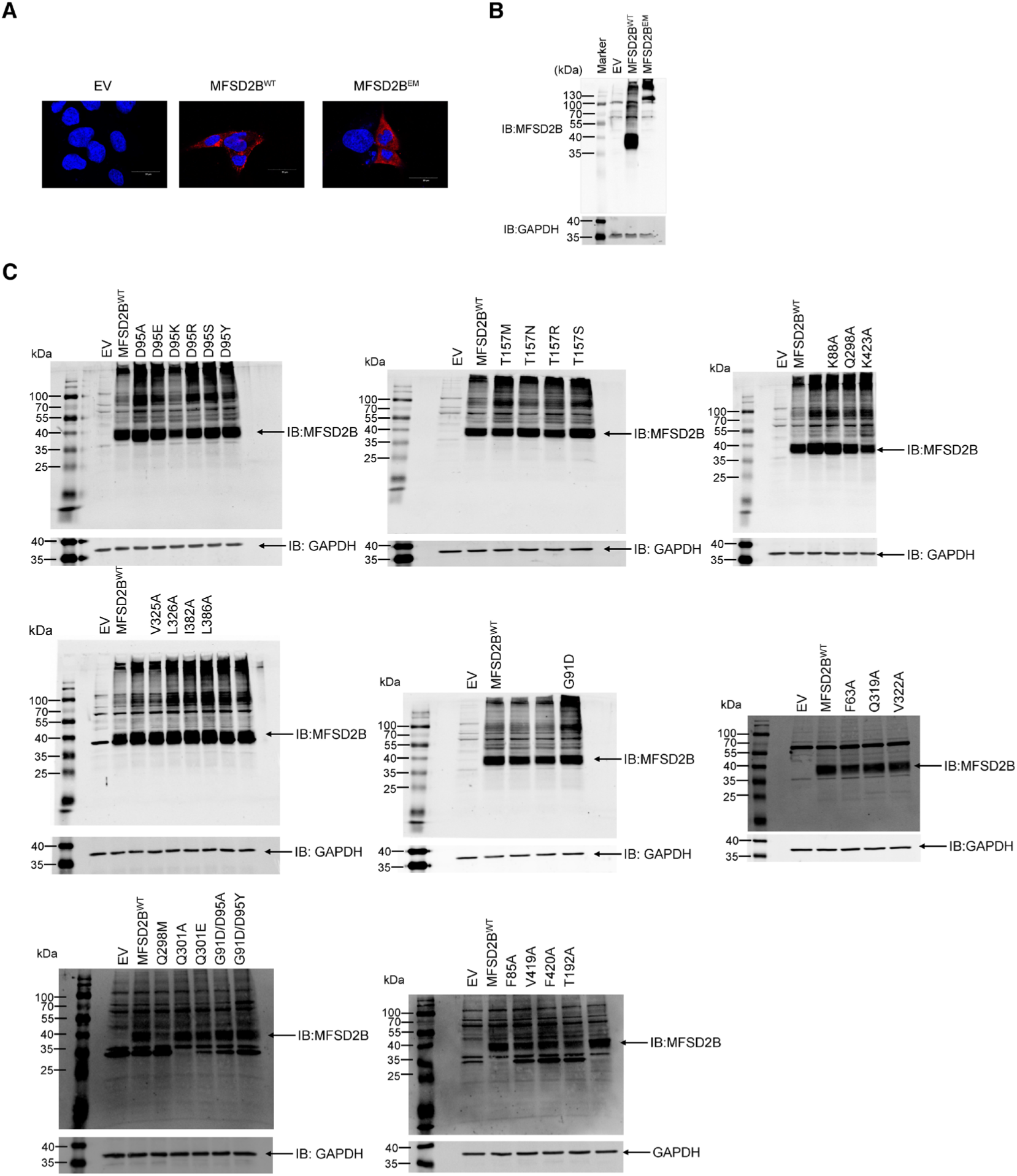
MFSD2B localization and expression. (A) Representative images showing the cellular localization of human MFSD2B^WT^ and MFSD2B^EM^. Cell nuclei were stained with Hoechst (blue), and MFSD2B proteins were stained with anti-MFSD2B and Alexa Fluor 555 (red). EV, empty vector–transfected cells. (B) Western blot analysis of MFSD2B expression levels. GAPDH served as a loading control. Protein molecular weight markers are indicated. (C) Expression levels of MFSD2B variants. GAPDH served as a loading control.

**Extended Data Fig. 2:**
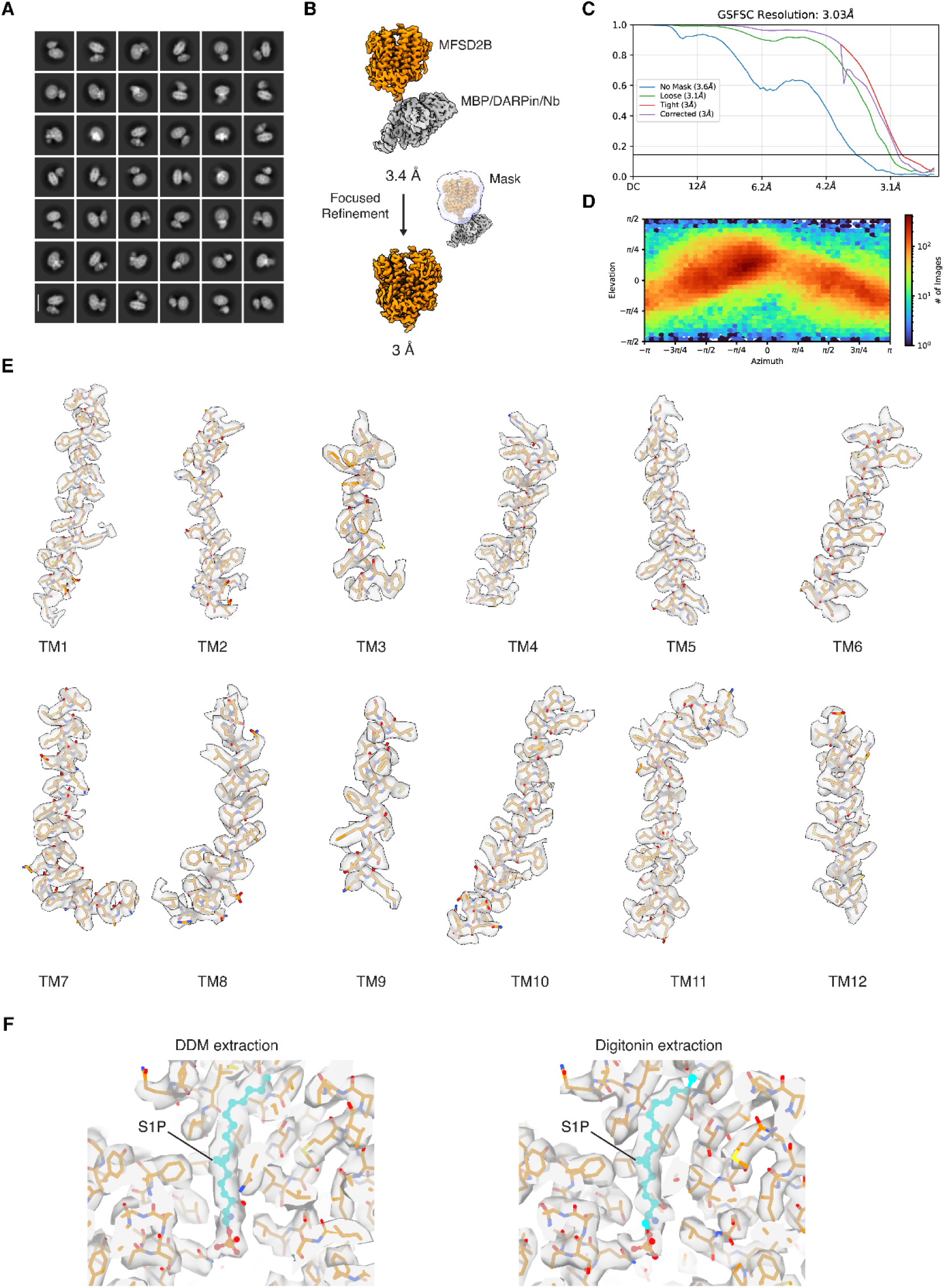
Cryo-EM analyses of MFSD2B in complex with S1P. (A) Representative 2D class averages of MFSD2B. (B) Final image processing step. MFSD2B colored in orange. (C) Fourier shell correlation (FSC) curves between two half maps. (D) Angular distribution of particles for the final 3D reconstruction. (E) Cryo-EM densities of the transmembrane helices. (F) Cryo-EM density around S1P-binding pocket. Left, MFSD2B extracted with DDM. Right, MFSD2B extracted with digitonin.

**Extended Data Fig. 3:**
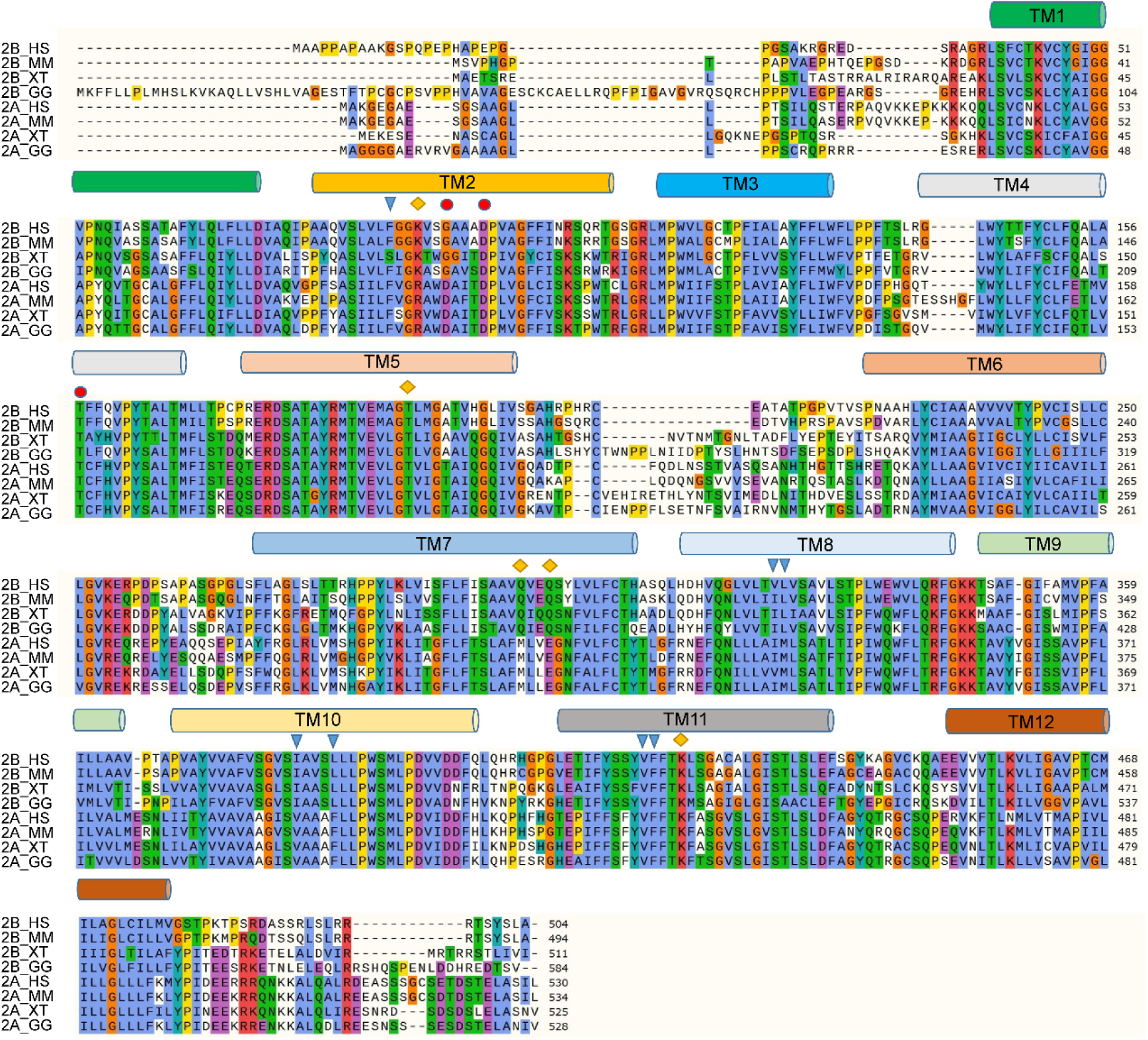
Sequence alignment of MFSD2B and MFSD2A homologs. Sequences of MFSD2B and MFSD2A from Homo sapiens (Hs), Mus musculus (Mm), Xenopus tropicalis (Xt), and Gallus gallus (Gg) are shown. Sequences were aligned using MUSCLE and visualized with the ClustalX color scheme. Residues discussed in this study are indicated. Rhombi denote headgroup-binding residues, triangles denote fatty-tail binding residues, and circles denote cation-sensing residues.

**Extended Data Fig. 4:**
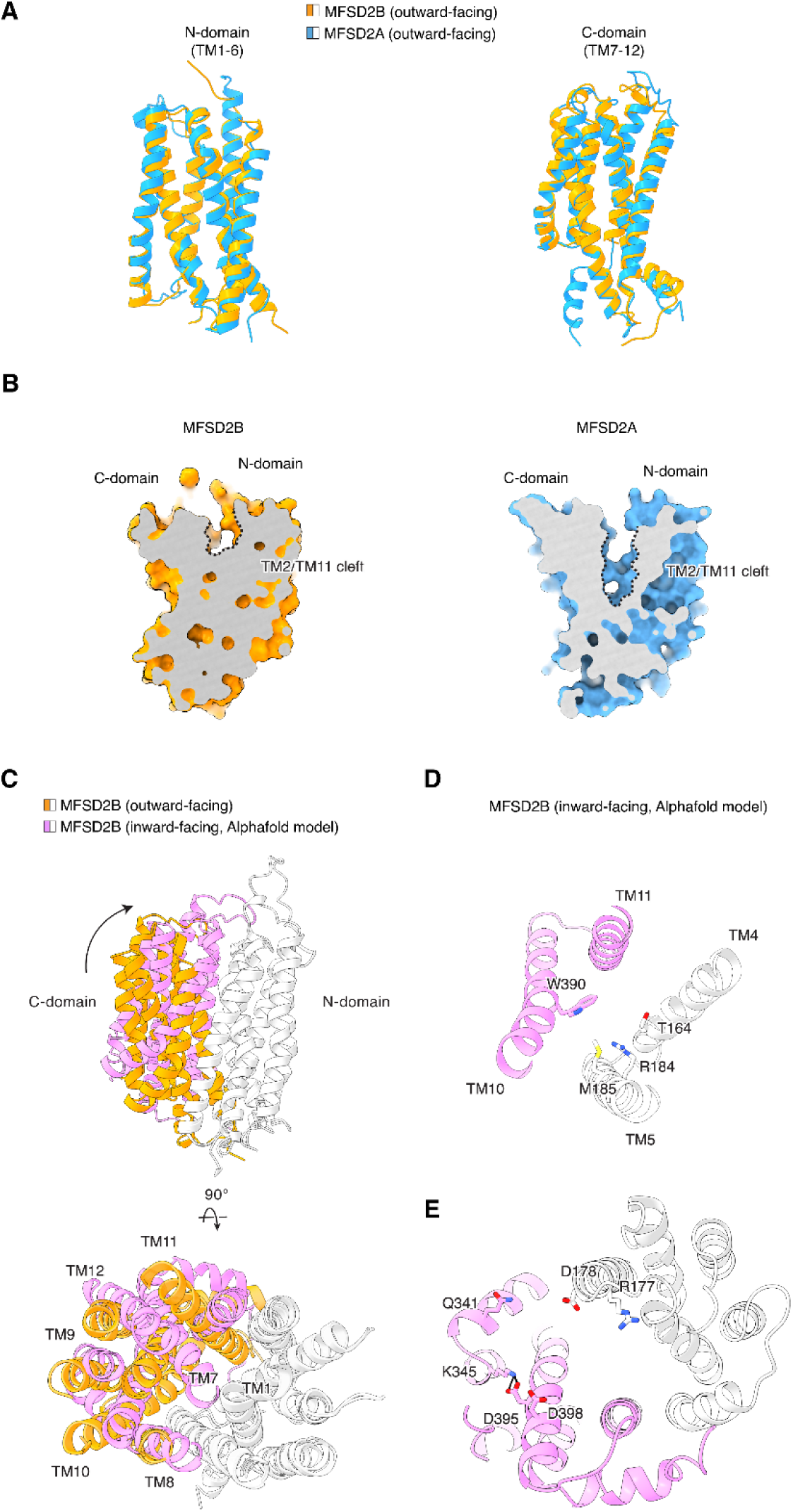
Structural analysis of MFSD2B. (A) Superimposition of the N-domain (left) and C-domain (right) of MFSD2B and MFSD2A. MFSD2B and MFSD2A are colored orange and blue, respectively. PDB code 7N98 for MFSD2A. (B) Cut-open view of MFSD2B and MFSD2A. (C) Superimposition of the experimental MFSD2B structure and the AlphaFold2 generated MFSD2B model. (D) Cytoplasmic gate of MFSD2B in a predicted inward-facing conformation. (E) Inter-domain interactions of MFSD2B in a predicted inward-facing conformation, viewed from the intracellular side.

**Extended Data Fig. 5:**
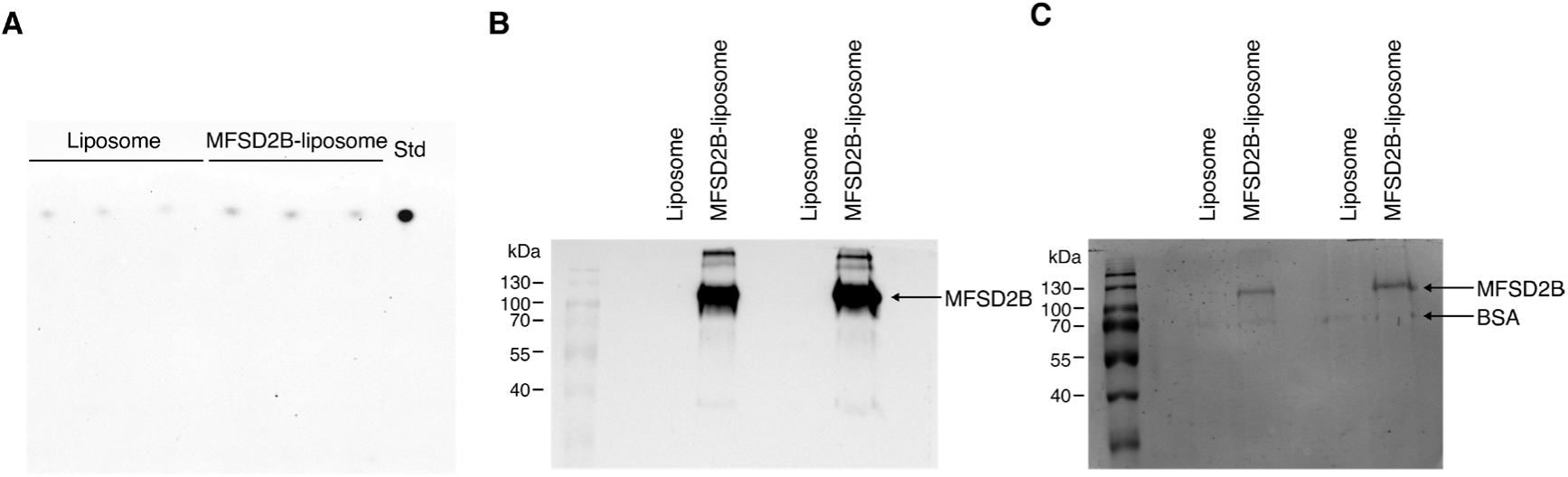
S1P transport in MFSD2B proteoliposomes. (A) NBD-S1P import by MFSD2B. (B) Western blot of MFSD2B proteoliposomes. (C) Ponceau staining of MFSD2B proteoliposomes.

**Extended Data Fig. 6:**
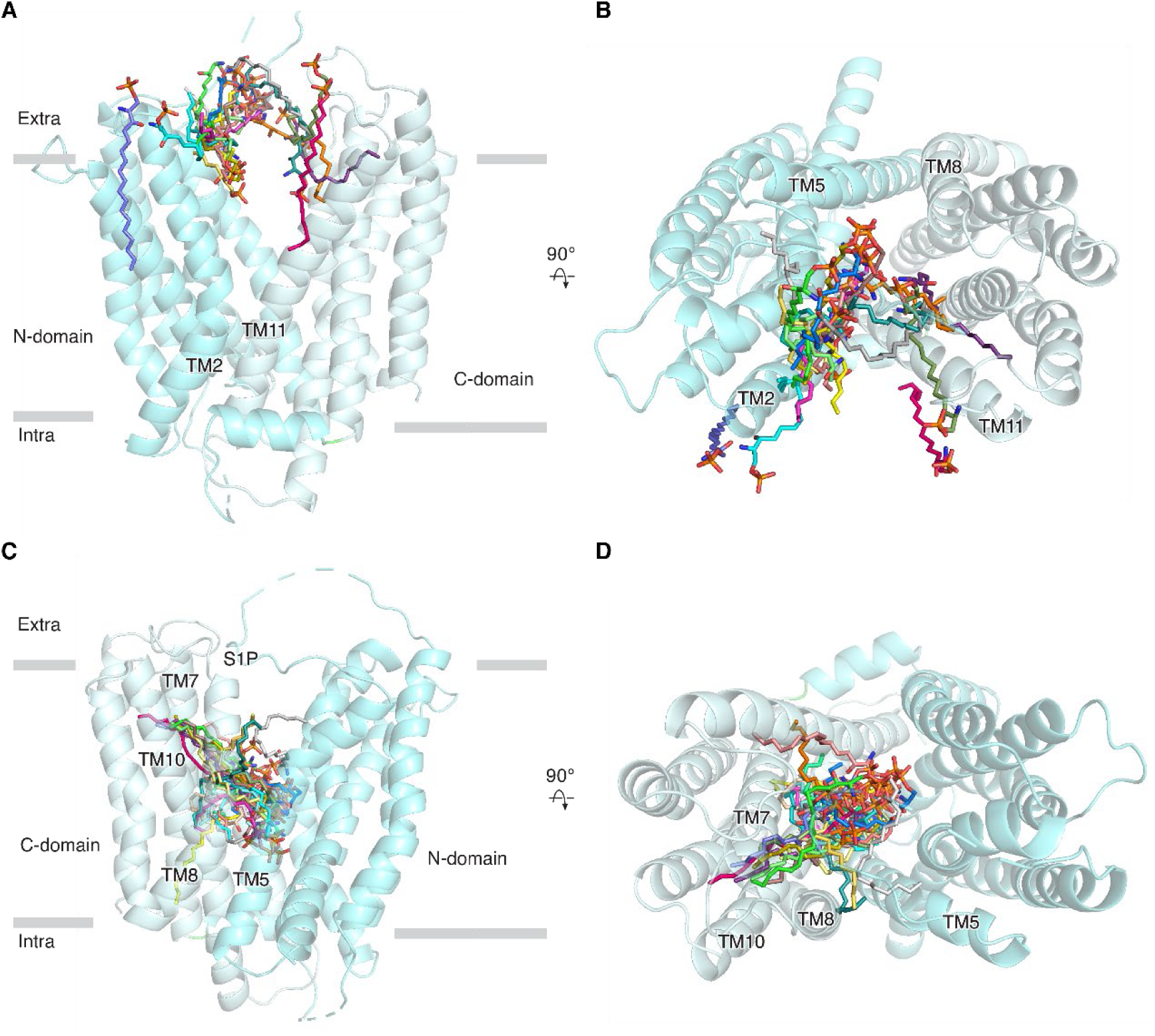
Molecular dynamics simulations of S1P entry into MFSD2B. (A) S1P orientations in MFSD2B during the 0–540 ns. S1P molecules are shown as sticks and colored according to different time points. (B) Same as panel A, but viewed from the extracellular side. (C) Conformations of S1P during the 1080–1800 ns time window. S1P molecules are shown as sticks. (D) Same as panel C, but viewed from the extracellular side.

**Extended Data Fig. 7:**
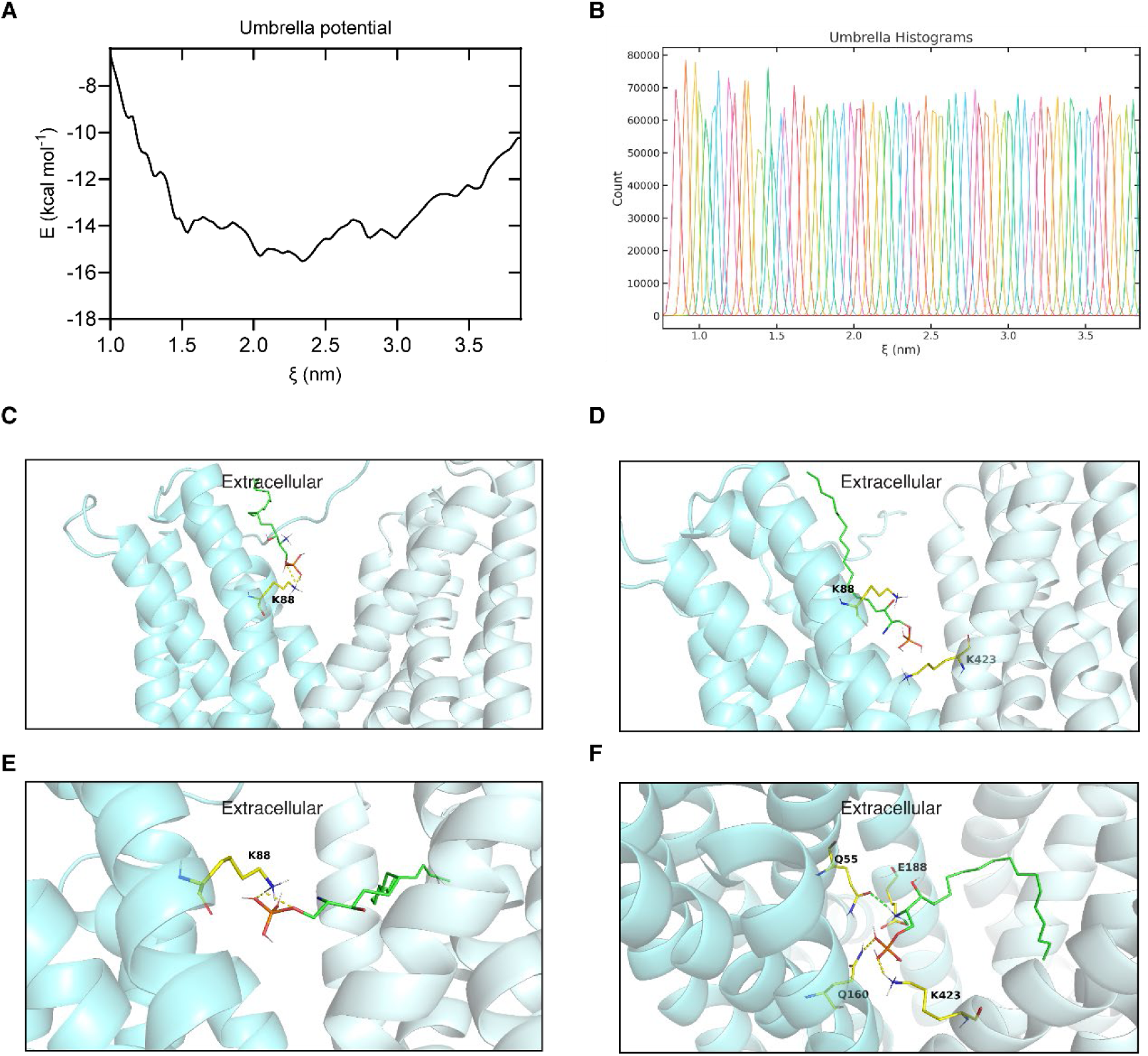
MF calculations. (A) All-atom PMF profile with four representative turning points along the trajectory (1, 510 ns; 2, 900 ns; 3, 1080 ns; 4, 1500 ns). (B) Umbrella histograms used for PMF calculations. (**C**–**F**) Snapshots of S1P conformations at turning points 1–4. S1P and key residues are shown as sticks. Salt bridges are colored in yellow and hydrogen bonds are colored in green. In panels C and E, S1P and K88 are highlighted; in panel D, S1P, K88, and K423 are shown; in panel F, S1P and surrounding residues are shown.

**Extended Data Fig. 8:**
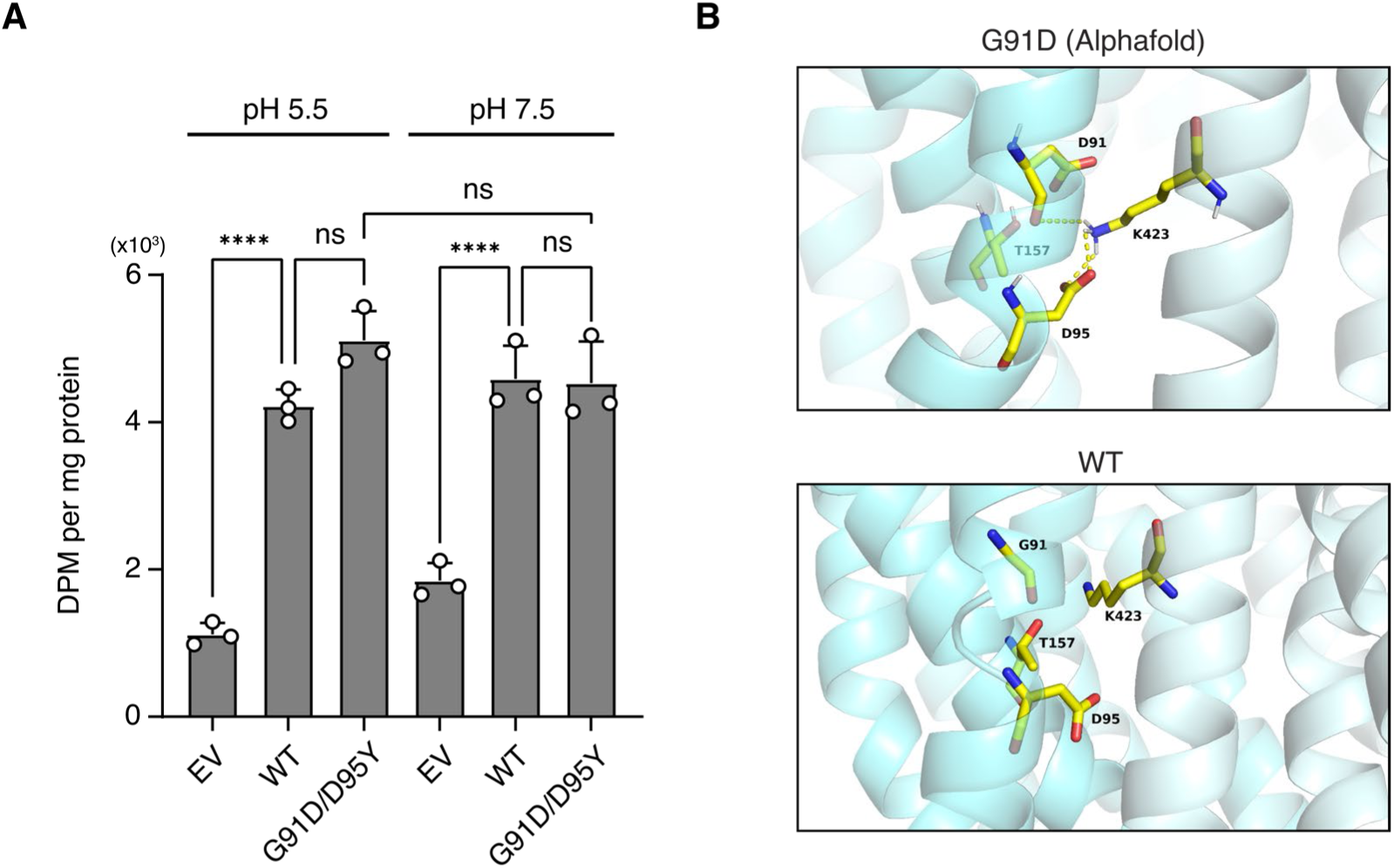
Analysis of MFSD2B coupling mechanism. (A) S1P export activity of MFSD2B^G91D/D95Y^. [³H]-S1P export at pH 5.5 or pH 7.5. Data are presented as mean ± SEM (n = 3 biological replicates). P values were calculated using one-way ANOVA followed by Dunnett’s multiple-comparison test. **** P < 0.0001; ns, not significant. (B) Structure of MFSD2B^G91D^ predicted by AlphaFold. Relevant residues are labeled, and salt bridges are shown in yellow.

**Extended Data Table 1:**
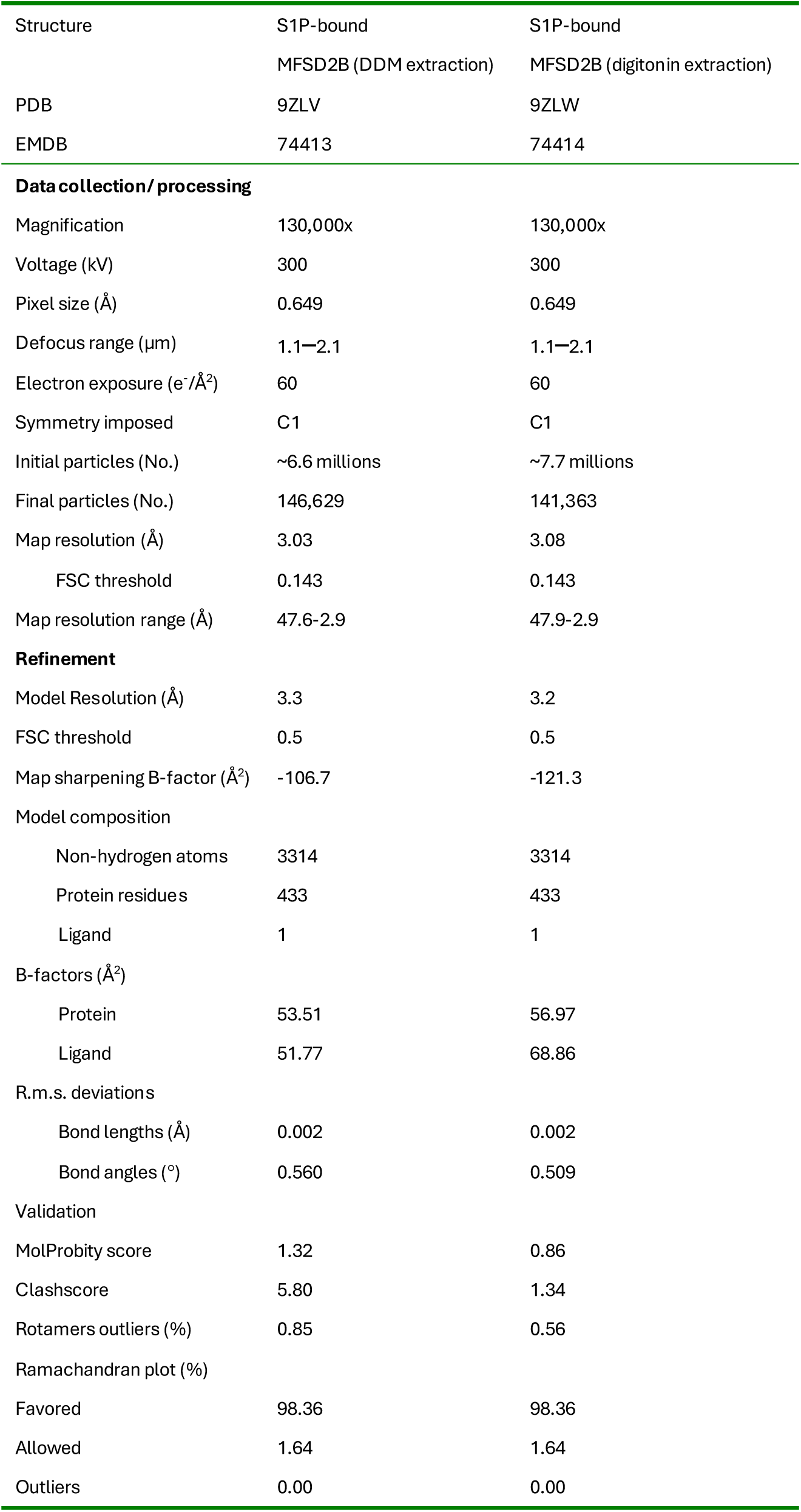
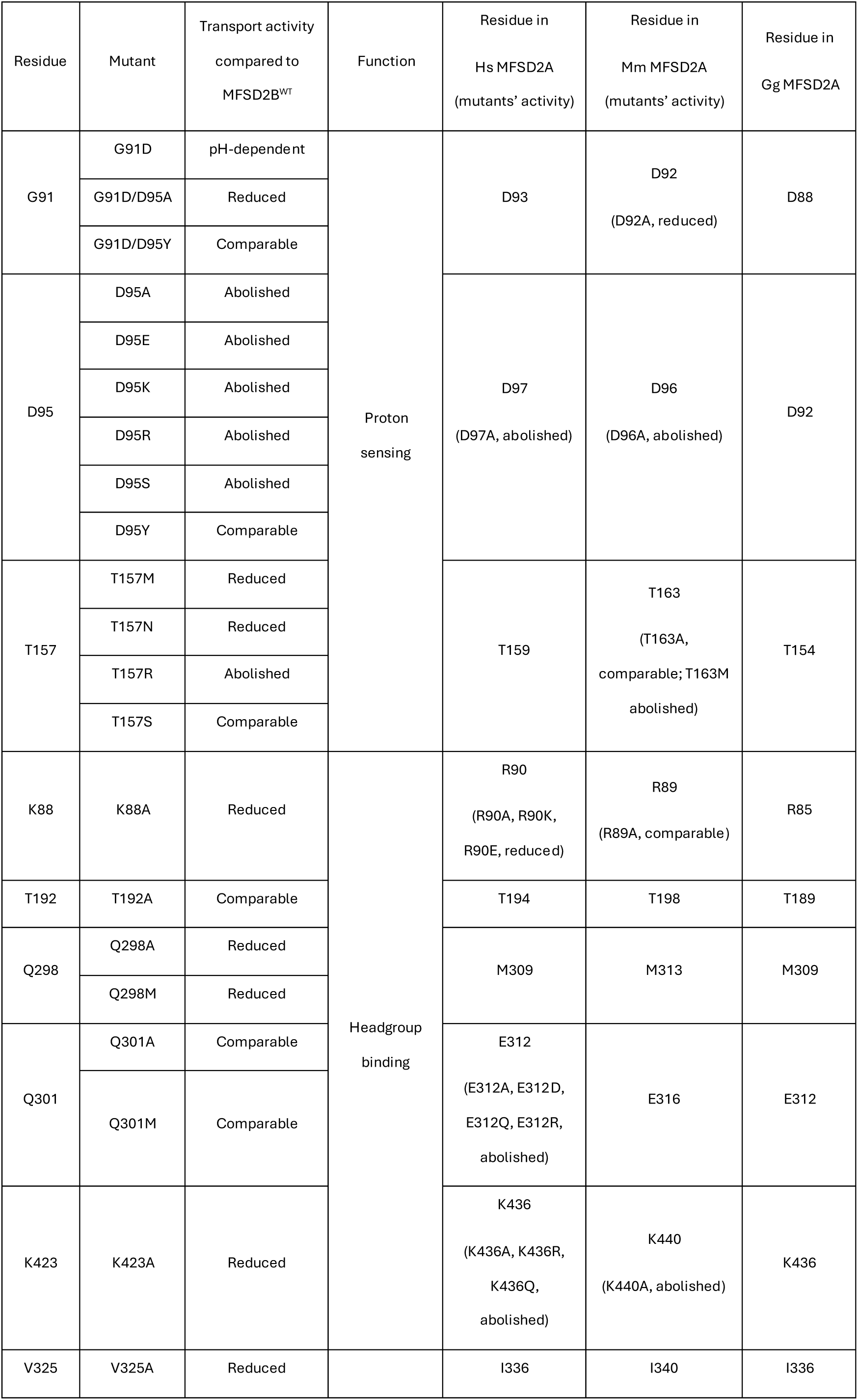
Cryo-EM data collection, refinement and validation statistics.

**Extended Data Table 2:**
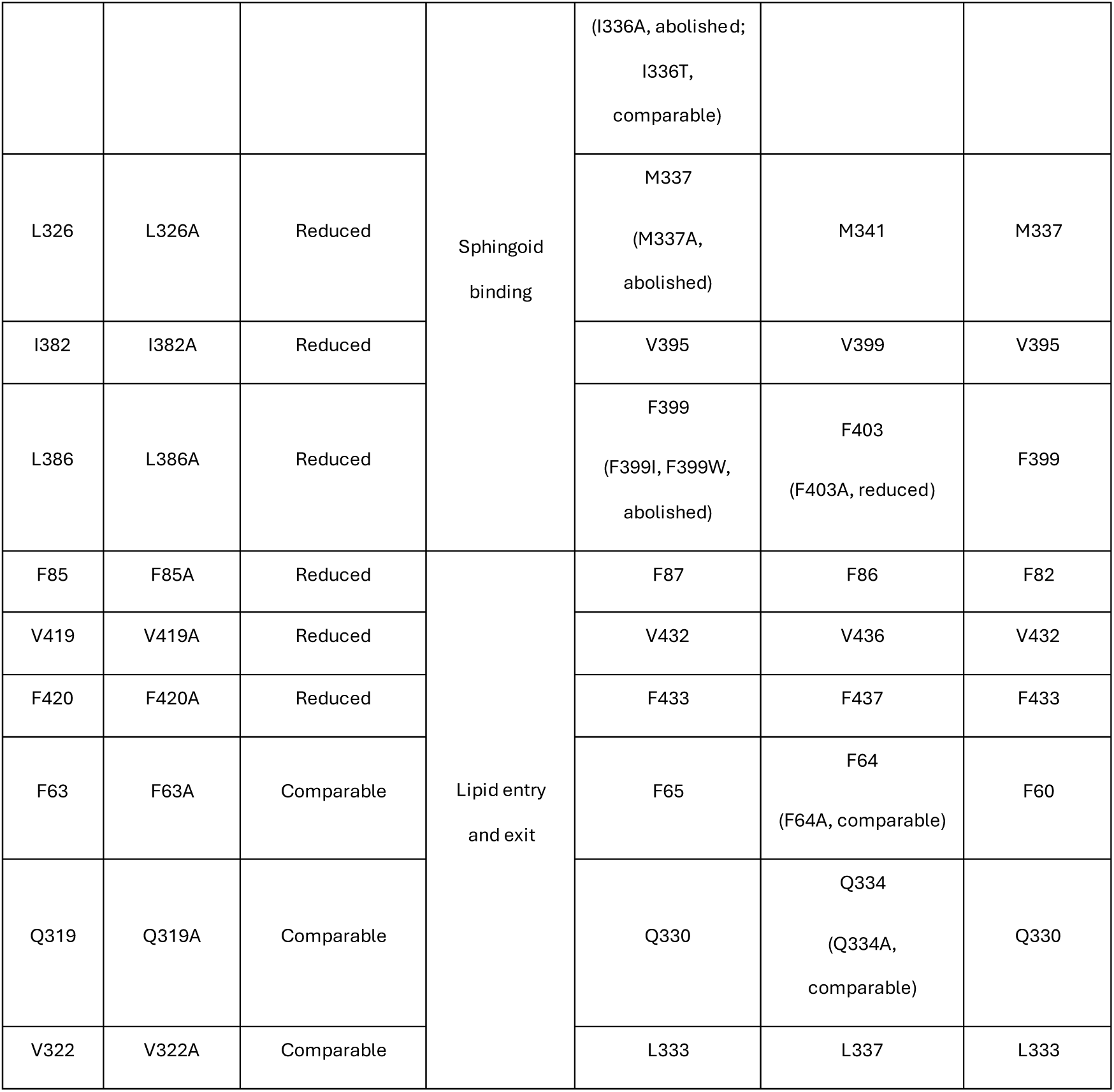
Transport activities of human MFSD2B mutants discussed in this study.

## References

1. Cartier, A. & Hla, T. Sphingosine 1-phosphate: Lipid signaling in pathology and therapy. Science 366, eaar5551 (2019).

2. Hla, T., Lee, M.J., Ancellin, N., Paik, J.H. & Kluk, M.J. Lysophospholipids--receptor revelations. Science 294, 1875–8 (2001).

3. Kuo, A. & Hla, T. Regulation of cellular and systemic sphingolipid homeostasis. Nat Rev Mol Cell Biol 25, 802–821 (2024).

4. Proia, R.L. & Hla, T. Emerging biology of sphingosine-1-phosphate: its role in pathogenesis and therapy. J Clin Invest 125, 1379–87 (2015).

5. Rosen, H., Stevens, R.C., Hanson, M., Roberts, E. & Oldstone, M.B. Sphingosine-1-phosphate and its receptors: structure, signaling, and influence. Annu Rev Biochem 82, 637–62 (2013).

6. Nguyen, T.Q. et al. Erythrocytes efficiently utilize exogenous sphingosines for S1P synthesis and export via Mfsd2b. J Biol Chem 296, 100201 (2021).

7. Spiegel, S., Maczis, M.A., Maceyka, M. & Milstien, S. New insights into functions of the sphingosine-1-phosphate transporter SPNS2. J Lipid Res 60, 484–489 (2019).

8. Spiegel, S. & Milstien, S. Functions of the multifaceted family of sphingosine kinases and some close relatives. J Biol Chem 282, 2125–9 (2007).

9. Kobayashi, N., Kobayashi, N., Yamaguchi, A. & Nishi, T. Characterization of the ATP-dependent sphingosine 1-phosphate transporter in rat erythrocytes. J Biol Chem 284, 21192–200 (2009).

10. Mitra, P. et al. Role of ABCC1 in export of sphingosine-1-phosphate from mast cells. Proc Natl Acad Sci U S A 103, 16394–9 (2006).

11. Kawahara, A. et al. The sphingolipid transporter spns2 functions in migration of zebrafish myocardial precursors. Science 323, 524–7 (2009).

12. Fukuhara, S. et al. The sphingosine-1-phosphate transporter Spns2 expressed on endothelial cells regulates lymphocyte trafficking in mice. J Clin Invest 122, 1416–26 (2012).

13. Osborne, N. et al. The spinster homolog, two of hearts, is required for sphingosine 1-phosphate signaling in zebrafish. Curr Biol 18, 1882–8 (2008).

14. Vu, T.M. et al. Mfsd2b is essential for the sphingosine-1-phosphate export in erythrocytes and platelets. Nature 550, 524–528 (2017).

15. Mendoza, A. et al. The transporter Spns2 is required for secretion of lymph but not plasma sphingosine-1-phosphate. Cell Rep 2, 1104–10 (2012).

16. Simmons, S. et al. High-endothelial cell-derived S1P regulates dendritic cell localization and vascular integrity in the lymph node. Elife 8(2019).

17. Hasan, Z. et al. Postnatal deletion of Spns2 prevents neuroinflammation without compromising blood vascular functions. Cell Mol Life Sci 79, 541 (2022).

18. Le, T.N.U. et al. Mfsd2b and Spns2 are essential for maintenance of blood vessels during development and in anaphylactic shock. Cell Rep 40, 111208 (2022).

19. Pappu, R. et al. Promotion of lymphocyte egress into blood and lymph by distinct sources of sphingosine-1-phosphate. Science 316, 295–8 (2007).

20. Venkataraman, K. et al. Vascular endothelium as a contributor of plasma sphingosine 1-phosphate. Circ Res 102, 669–76 (2008).

21. Chandrakanthan, M. et al. Deletion of Mfsd2b impairs thrombotic functions of platelets. Nat Commun 12, 2286 (2021).

22. Duse, D.A. et al. Deficiency of the sphingosine-1-phosphate (S1P) transporter Mfsd2b protects the heart against hypertension-induced cardiac remodeling by suppressing the L-type-Ca(2+) channel. Basic Res Cardiol 119, 853–868 (2024).

23. Ha, H.T., et al. Lack of SPNS1 results in accumulation of lysolipids and lysosomal storage disease in mouse models. JCI Insight 9, e175462 (2024).

24. He, M. et al. Spns1 is a lysophospholipid transporter mediating lysosomal phospholipid salvage. Proc Natl Acad Sci U S A 119, e2210353119 (2022).

25. Nguyen, L.N. et al. Mfsd2a is a transporter for the essential omega-3 fatty acid docosahexaenoic acid. Nature 509, 503–6 (2014).

26. Bergman, S., Cater, R.J., Plante, A., Mancia, F. & Khelashvili, G. Substrate binding-induced conformational transitions in the omega-3 fatty acid transporter MFSD2A. Nat Commun 14, 3391 (2023).

27. Cater, R.J. et al. Structural basis of omega-3 fatty acid transport across the blood-brain barrier. Nature 595, 315–319 (2021).

28. Chen, H. et al. Structural and functional insights into Spns2-mediated transport of sphingosine-1-phosphate. Cell 186, 2644–2655 e16 (2023).

29. Chen, H. et al. Molecular basis of Spns1-mediated lysophospholipid transport from the lysosome. Proc Natl Acad Sci U S A 122, e2409596121 (2025).

30. Duan, Y. et al. Structural basis of Sphingosine-1-phosphate transport via human SPNS2. Cell Res 34, 177–180 (2024).

31. Li, H.Z. et al. Transport and inhibition of the sphingosine-1-phosphate exporter SPNS2. Nat Commun 16, 721 (2025).

32. Martinez-Molledo, M., Nji, E. & Reyes, N. Structural insights into the lysophospholipid brain uptake mechanism and its inhibition by syncytin-2. Nat Struct Mol Biol 29, 604–612 (2022).

33. Nguyen, C. et al. Lipid flipping in the omega-3 fatty-acid transporter. Nat Commun 14, 2571 (2023).

34. Pang, B. et al. Molecular basis of Spns2-facilitated sphingosine-1-phosphate transport. Cell Res 34, 173–176 (2024).

35. Wood, C.A.P. et al. Structure and mechanism of blood-brain-barrier lipid transporter MFSD2A. Nature 596, 444–448 (2021).

36. Pidathala, S. et al. Mechanisms of neurotransmitter transport and drug inhibition in human VMAT2. Nature 623, 1086–1092 (2023).

37. Dai, Y. & Lee, C.H. Transport mechanism and structural pharmacology of human urate transporter URAT1. Cell Res 34, 776–787 (2024).

38. Pidathala, S. et al. Structural pharmacology of SV2A reveals an allosteric modulation mechanism in the major facilitator superfamily. Nat Commun 16, 10748 (2025).

39. Jumper, J. et al. Highly accurate protein structure prediction with AlphaFold. Nature 596, 583–589 (2021).

40. Goto, H., Miyamoto, M. & Kihara, A. Direct uptake of sphingosine-1-phosphate independent of phospholipid phosphatases. J Biol Chem 296, 100605 (2021).

41. Trzesniak, D., Kunz, A.P.E. & van Gunsteren, W.F. A Comparison of Methods to Compute the Potential of Mean Force. ChemPhysChem 8, 162–169 (2006).

42. Binz, H.K. et al. High-affinity binders selected from designed ankyrin repeat protein libraries. Nat Biotechnol 22, 575–82 (2004).

43. Goehring, A. et al. Screening and large-scale expression of membrane proteins in mammalian cells for structural studies. Nat Protoc 9, 2574–85 (2014).

44. Kirchhofer, A. et al. Modulation of protein properties in living cells using nanobodies. Nat Struct Mol Biol 17, 133–8 (2010).

45. Zimmermann, I. et al. Synthetic single domain antibodies for the conformational trapping of membrane proteins. Elife 7(2018).

46. Punjani, A., Rubinstein, J.L., Fleet, D.J. & Brubaker, M.A. cryoSPARC: algorithms for rapid unsupervised cryo-EM structure determination. Nat Methods 14, 290–296 (2017).

47. Bepler, T. et al. Positive-unlabeled convolutional neural networks for particle picking in cryo-electron micrographs. Nat Methods 16, 1153–1160 (2019).

48. Punjani, A., Zhang, H. & Fleet, D.J. Non-uniform refinement: adaptive regularization improves single-particle cryo-EM reconstruction. Nat Methods 17, 1214–1221 (2020).

49. Pettersen, E.F. et al. UCSF Chimera--a visualization system for exploratory research and analysis. J Comput Chem 25, 1605–12 (2004).

50. Emsley, P., Lohkamp, B., Scott, W.G. & Cowtan, K. Features and development of Coot. Acta Crystallogr D Biol Crystallogr 66, 486–501 (2010).

51. Croll, T.I. ISOLDE: a physically realistic environment for model building into low-resolution electron-density maps. Acta Crystallogr D Struct Biol 74, 519–530 (2018).

52. Afonine, P.V. et al. Real-space refinement in PHENIX for cryo-EM and crystallography. Acta Crystallogr D Struct Biol 74, 531–544 (2018).

53. Chen, V.B. et al. MolProbity: all-atom structure validation for macromolecular crystallography. Acta Crystallogr D Biol Crystallogr 66, 12–21 (2010).

54. Meng, E.C. et al. UCSF ChimeraX: Tools for structure building and analysis. Protein Sci 32, e4792 (2023).

55. Abraham, M.J. et al. GROMACS: High performance molecular simulations through multi-level parallelism from laptops to supercomputers. SoftwareX 1-2, 19-25 (2015).

56. Jo, S., Kim, T., Iyer, V.G. & Im, W. CHARMM-GUI: a web-based graphical user interface for CHARMM. J Comput Chem 29, 1859–65 (2008).

57. Huang, J. et al. CHARMM36m: an improved force field for folded and intrinsically disordered proteins. Nat Methods 14, 71–73 (2017).

58. Essmann, U. et al. A smooth particle mesh Ewald method. J Chem Phys 103, 8577–8593 (1995).

59. Hess, B., Bekker, H., Berendsen, H.J.C. & Fraaije, J.G.E.M. LINCS: A linear constraint solver for molecular simulations. J Comput Chem 18, 1463–1472 (1997).

60. Berendsen, H.J.C., Postma, J.P.M., van Gunsteren, W.F., DiNola, A. & Haak, J.R. Molecular dynamics with coupling to an external bath. J Chem Phys 81, 3684–3690 (1984).

61. Bussi, G., Donadio, D. & Parrinello, M. Canonical sampling through velocity rescaling. J Chem Phys 126, 014101 (2007).

62. Kumar, S., Bouzida, D., Swendsen, R.H., Kollman, P.A. & Rosenberg, J.M. THE weighted histogram analysis method for free - energy calculations on biomolecules. I. The method. J Comput Chem 13, 1011–1021 (1992).

